# ImPaqT - A Golden Gate-based Toolkit for Zebrafish Transgenesis

**DOI:** 10.1101/2024.08.02.606358

**Authors:** Saskia Hurst, Christiane Dimmler, Mark R. Cronan

## Abstract

Transgenic animals continue to play an essential role in many aspects of zebrafish research, including the development of disease models. The most widely used system for zebrafish transgenesis is the Tol2 transposon system. Here, we have developed ImPaqT (**Im**munological toolkit for **Paq**CI-based Golden Gate Assembly of Tol2 **T**ransgenes), a new Tol2-based transgenesis system that utilizes Golden Gate assembly to facilitate the production of transgenic zebrafish lines. This system allows for rapid assembly of multiple fragments into a single transgene, facile swapping of individual sequences to generate new transgenes and an easy cloning workflow to incorporate new genetic elements into the existing kit. Within this toolkit framework, we have generated a number of reagents to enable gene expression within immune and non-immune cell types, an array of best-in-class fluorescent proteins to visualize cell populations and transgenes as well as tools to simplify genetic manipulation, purification and ablation of targeted cells. Unlike recombination-based systems, the Golden Gate approach is also expandable, allowing the incorporation of complex designs such as multi-fragment promoters within the established modular framework of ImPaqT. Here, we demonstrate the function of this new system by generating a number of novel transgenic immune reporter lines. While our toolkit is focused on the immune system as an emerging area of study within zebrafish research, the ImPaqT approach can be broadly adapted to the construction of almost any zebrafish transgene, offering new tools for the generation of transgenes within the zebrafish community.

## Introduction

The zebrafish (*Danio rerio*) has emerged as a powerful model organism and is increasingly used in various research fields and disciplines. Zebrafish share 70% of their genome and more than 80% of disease genes with humans (Howe et al. 2013). This has led to the development of many distinct zebrafish disease models (Santoriello and Zon 2012) including models of hematopoietic and neurodegenerative diseases, cardiovascular diseases (Bowley et al. 2022) such as atherosclerosis (Tang et al. 2021), cancer (Liu and Leach 2011; Russo et al. 2022), and tuberculosis (Bouz and Al Hasawi 2018; Varela and Meijer 2022), among others. Transgenic animals continue to play a key role for many experimental strategies, including study of gene function, genetic screens, and development of disease models. Transgenic zebrafish lines have been widely generated by applying the Tol2 transposon system with a transgenesis rate of 50% (Kawakami et al. 2004). In this system, a transposon-donor plasmid containing a DNA sequence of interest flanked by *Tol2* sites and synthetic mRNA encoding the Tol2 transposase are introduced into the single cell of fertilized eggs by microinjection. Upon translation, the transposase protein integrates the *Tol2* flanked gene fragment randomly into the genome through a cut-and-paste mechanism without causing rearrangement or modifications at the target site (Kawakami and Shima 1999; Kawakami et al. 2000, 2004; Kawakami 2007).

A number of approaches have previously been used to generate Tol2 transgenesis constructs. Traditionally, multi-step restriction enzyme and ligase based subcloning was used. However, this has given way to many advanced cloning approaches, including Gateway cloning (Hartley et al. 2000; Kwan et al. 2007; Katzen 2007), Gibson Assembly (Gibson et al. 2009), In-Fusion cloning (Zhu et al. 2007; Sleight et al. 2010) and Golden Gate cloning, which have been used to speed up and simplify the procedure of transgene assembly. Each of these approaches have different advantages and disadvantages. Gateway cloning allows for the establishment and reuse of a library of fragments (e.g. promoters, genes and fluorescent proteins) but requires the use of lambda phage-derived *att* sequences to facilitate insertion of the desired equences. These *att* sequences result in the addition of ∼20-25 bp of additional sequence in between each element and expansion of the number of fragments to be assembled can require recloning of the sequence with new *att* ends (Katzen 2007; Engler et al. 2008; Sone et al. 2008). By contrast, Gibson and In-Fusion cloning allow for joining of short (15-20 bp) regions of homology, enabling the seamless cloning of many transgenes. However, the requirement for homology for seamless cloning generally means that individual components have to be adapted to upstream and downstream elements to generate new transgenes and have an upper limit of approximately 5-6 fragments with currently available kits. Golden Gate cloning utilizes type IIS restriction enzymes to cleave and join multiple fragments in a one pot reaction (Padgett and Sorge 1996; Engler et al. 2008). It balances the tradeoffs of the previous cloning systems, requiring inclusion of only a short 4 base pair sequence, while enabling researchers to generate a library of vectors that can be combinatorially assembled to generate distinct transgenes. Large numbers of DNA fragments can also be assembled by Golden Gate with reports of assembly of up to 52 fragments in a single reaction (Pryor et al. 2022). Despite the advantages of Golden Gate cloning, this approach has not been widely applied in zebrafish transgene assembly. Only a few examples of this approach have been described for Tol2 transgenesis including the recently described GoldenFish toolkit which uses Golden Gate cloning to gain more flexibility in transgene design (Jiang et al. 2022) and the Golden Gateway tool (Kirchmaier et al. 2013) which combines the well-established cloning tools Gateway and Golden Gate cloning to simplify creation of new components for Gateway assembly of transgenes.

Here, we have sought to apply Golden Gate cloning approaches to create a toolkit for zebrafish transgenesis. We have developed ImPaqT, a new **Im**munological toolkit for **Paq**CI-based Golden Gate Assembly of Tol2 **T**ransgenes, to facilitate the production of transgenic animals. Studies of immune function have been of growing interest in the zebrafish community due to the close genetic similarity of the immune systems of humans and zebrafish (Howe et al. 2013), but there are few compiled resources for these cell types. ImPaqT provides a collection of commonly used immunological promoters targeting distinct lineages of vertebrate immunity, genetic tools and fluorophores supporting flexible generation of transgenes for immunological research. This toolkit utilizes Golden Gate cloning due to the advantages of this technique and incorporates a newly described type IIS restriction enzyme, PaqCI, with a 7 base recognition sequence to reduce the frequency of off target cutting (from 1 in 4,096 bases (4^6^) for a 6-base cutter to 1 in 16,384 bases (4^7^), assuming random cutting). We designed the kit to have consistent overhang sequences for each position in the backbone (e.g. 5’, middle, and 3’ element) in order to make the toolkit modular and simplify the construction of additional transgenes. As proof-of-concept, we show that this toolkit can facilitate the production of expression constructs and we demonstrate the generation of transgenic animals using the Tol2 transgenesis with these constructs. ImPaqT is modular, enabling easy swapping of gene sequences, easily expandable, allowing the introduction of multiple new genetic elements if needed, and relies only on the use of short (4 bp) overlaps that can also be designed to be scarless if desired. To establish an immunological toolkit that will be broadly useful to zebrafish researchers, we have generated a variety of immunological and non-immunological promoters, including *mfap4* and *mpeg1.1* (macrophages)*, irg1* (macrophages, induced by infection)*, lyz* (neutrophils), *lck* (T cells), *ubb* (ubiquitious) and *hsp70* (heat shock inducible), tools for genetic recombination (*icre*), cell killing (*kid, nitroreductase)* and cell isolation tools (*LNGFR*), and an array of fluorescence proteins including *mTurquoise2*, *tdStayGold*, *mCitrine*, *mNeonGreen*, *dLanYFP* and *tdTomato* that can be used to rapidly develop new transgenes and new transgenic lines.

## Results

### PaqCI-based Golden Gate cloning enables the construction of Tol2 expression vectors

We sought to create a toolkit for immune transgene construction for the zebrafish community. Ultimately, we decided to design this toolkit to use Golden Gate cloning, given the short regions of homology that this approach required, the ability to structure this toolkit in a modular fashion and the expandability of the Golden Gate cloning approach, enabling incorporation of additional genetic elements as needed. Golden Gate assembly uses type IIS restriction enzymes to generate a set of unique overhangs that are used to join fragments in a desired order. To create a system for modular cloning of transgenes, we chose consistent 4-base overhang sequences for each of the three elements such that DNA fragments amplified with these overhangs can be cloned in the same position in the final construct, allowing these elements to be used combinatorially. The individual overhang sequences were chosen using NEBridge Ligase Fidelity Viewer (https://ligasefidelity.neb.com/viewset/run.cgi), which enables the selection of efficient joining ends with minimal off-target ligation based on high throughput, combinatorial tests of all 4-base overhangs (Pryor et al. 2020; Potapov et al. 2018). Using this tool, we selected ends predicted to have 1% or less off-target ligation with the other ends used in the toolkit. As zebrafish transgenes generally consist of a cell-type specific promoter, a gene of interest, and a fluorescent marker, we structured our toolkit to have three elements that would ultimately be joined within a backbone vector possessing Tol2 repeats for transgenesis. These elements were named based on their location within the final construct - 5’ element (5E) for the first position, a middle element (ME) and a 3’ element (3E) for the final position within the backbone.

To generate the overhangs required to join individual components for Golden Gate assembly, we chose the 7-base cutter PaqCI to limit off-target cutting of inserts used in assembly. Searches of the zebrafish genome revealed that PaqCI cuts once every ∼17 kb on average (∼82600 sites in the GRCz11 zebrafish genome assembly), while the 6-base cutters BsaI and BsmBI, commonly used for Golden Gate, cut every ∼9 kb (∼151800 sites) or ∼13 kb (∼107200 sites). Genes and regions of interest were generated by PCR amplification using primers containing the 7-base PaqCI recognition sequence along with a spacer consisting of any four nucleotides that is excised from the final construct, and a 4-base overhang sequence defined by the location of the sequence of interest in the final construct (Fig. S1). These PCR products were cloned into vectors and sequence verified to generate a stable library of constructs for transgene assembly. Cleavage with PaqCI leads to the creation of specific 4-base overhangs for each position at their 5’ and 3’ end enabling proper assembly and single-step ligation into a backbone vector for transgene generation. These backbone vectors possess inverted *Tol2* repeats for transgenesis surrounding a PaqCI-flanked LacZ gene for blue-white screening to identify backbone constructs containing transgene components. A simplified schematic for the assembly using one insert is illustrated in Fig. 1. As we demonstrated below, we have found that this approach allows for efficient, combinatorial transgene generation.

**Figure 1:**
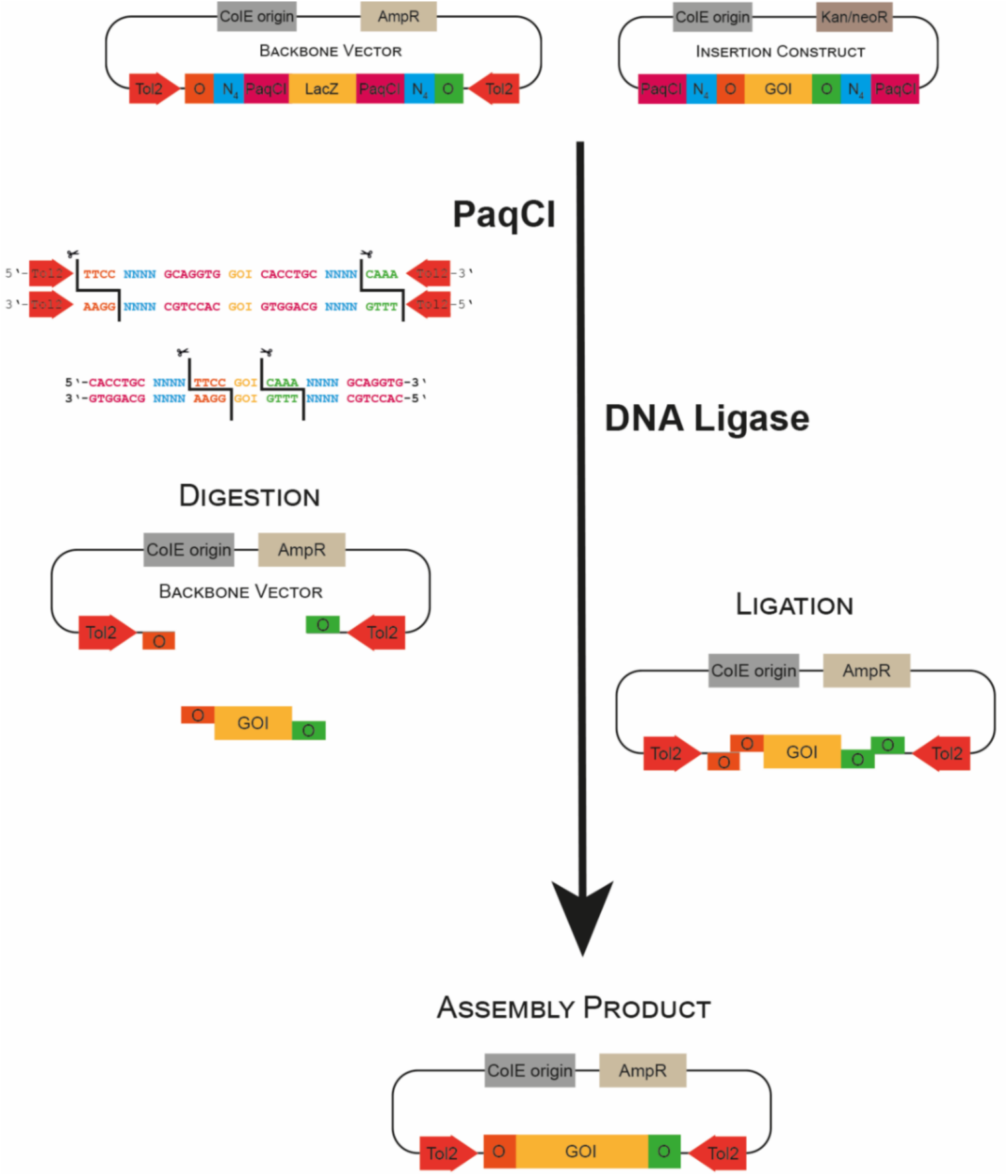
Basic principles of Golden Gate cloning for Tol2 Transgenesis Schematic of a Golden Gate cloning reaction using one insert. The insertion construct and backbone vector are assembled in a single-tube reaction using PaqCI and ligase. PaqCI is a type IIS restriction enzyme. Its recognition site and cleavage are indicated. The font color corresponds to the color of the respective feature indicated in the cartoon. The backbone vector consists of Tol2 sites, PaqCI recognition sites (PaqCI) along with 4-base spacer sequences (N_4_) and two overhangs (O). The insertion construct includes the gene of interest (GOI) flanked by PaqCI, N_4_ and the same two overhangs. Assembly involves digestion of backbone and insertion constructs by PaqCI producing matching overhangs and ligation through the DNA ligase forming the final assembly product.

### ImPaqT facilitates the flexibility of transgenes design and the generation of transgenic zebrafish

Having established this Golden Gate cloning-based technique for the assembly of Tol2 transposon vectors, we sought to create an array of constructs that would be beneficial to the immunology community to generate transgenic animals. To these ends, we have generated constructs for expression in commonly studied zebrafish immune populations, including T cells (*lck*) (Langenau et al. 2004), neutrophils (*lyz)* (C. Hall et al. 2007) and macrophages (*mfap4*, *mpeg1.1* and the infection-inducible *irg1*) (Walton et al. 2015; Ellett et al. 2011; C. J. Hall et al. 2013). To enable lineage tracing approaches for labeling of hematopoietic derived cells, we also included the hematopoietic stem cell (HSC)-specific promoter *runx1 +23* (Tamplin et al. 2015) as well as tools for broader expression including the ubiquitious promoter *ubiquitin B* (*ubb*) (Mosimann et al. 2011), and the temperature inducible heat shock protein 70-like promoter (*hsp70*) (Shoji et al. 1998; Halloran et al. 2000; Kwan et al. 2007). We have also integrated a number of tools to allow for manipulation of these immune populations, including tools for genetic manipulation (*icre* (Shimshek et al. 2002)), for rapid purification of cell populations (truncated human *LNGFR* (Wattrus and Zon 2020)), expression CRISPR gRNAs and other small RNAs (zebrafish U6 promoter (Zhou et al. 2018)) and cell-specific ablation (*nitroreductase* (Curado et al. 2007, 2008) and *kid* (Labbaf et al. 2022)). To visualize labeled cell populations and genes, we provide a diverse array of best-in-class fluorescent proteins in a range of colors to facilitate simultaneous imaging of distinct cell types. These include *tdStayGold* (Hirano et al. 2022), *mTurquoise2* (Goedhart et al. 2012), *tdTomato* (Shaner et al. 2004), m*Citrine* (Zacharias et al. 2002)*, mNeonGreen* (Shaner et al. 2013) and *dLanYFP* (Shaner et al. 2013). A summary of all available constructs is shown in Table 1. ImPaqT is also designed to be easily expandable through simple cloning approaches.

**Table 1:**
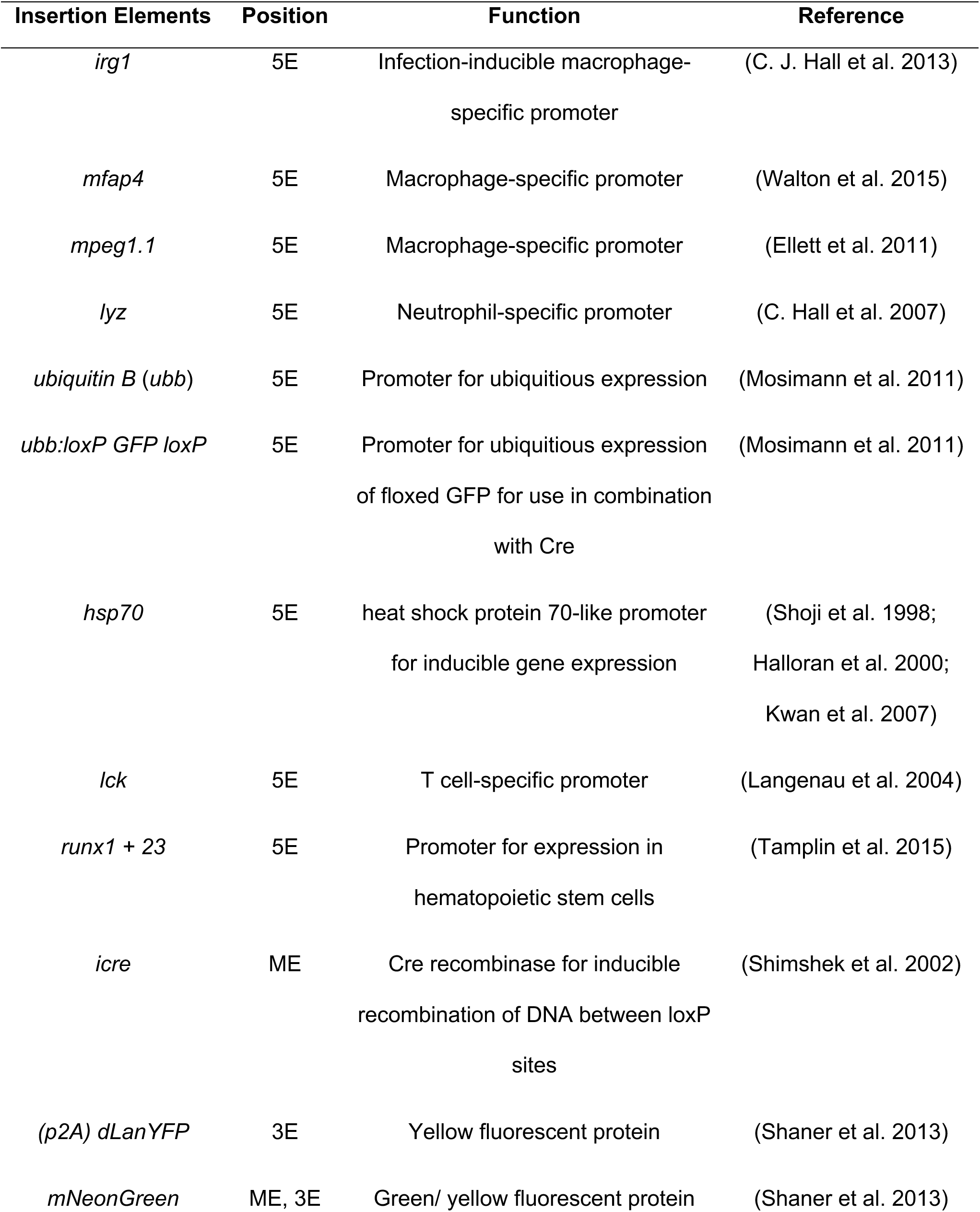
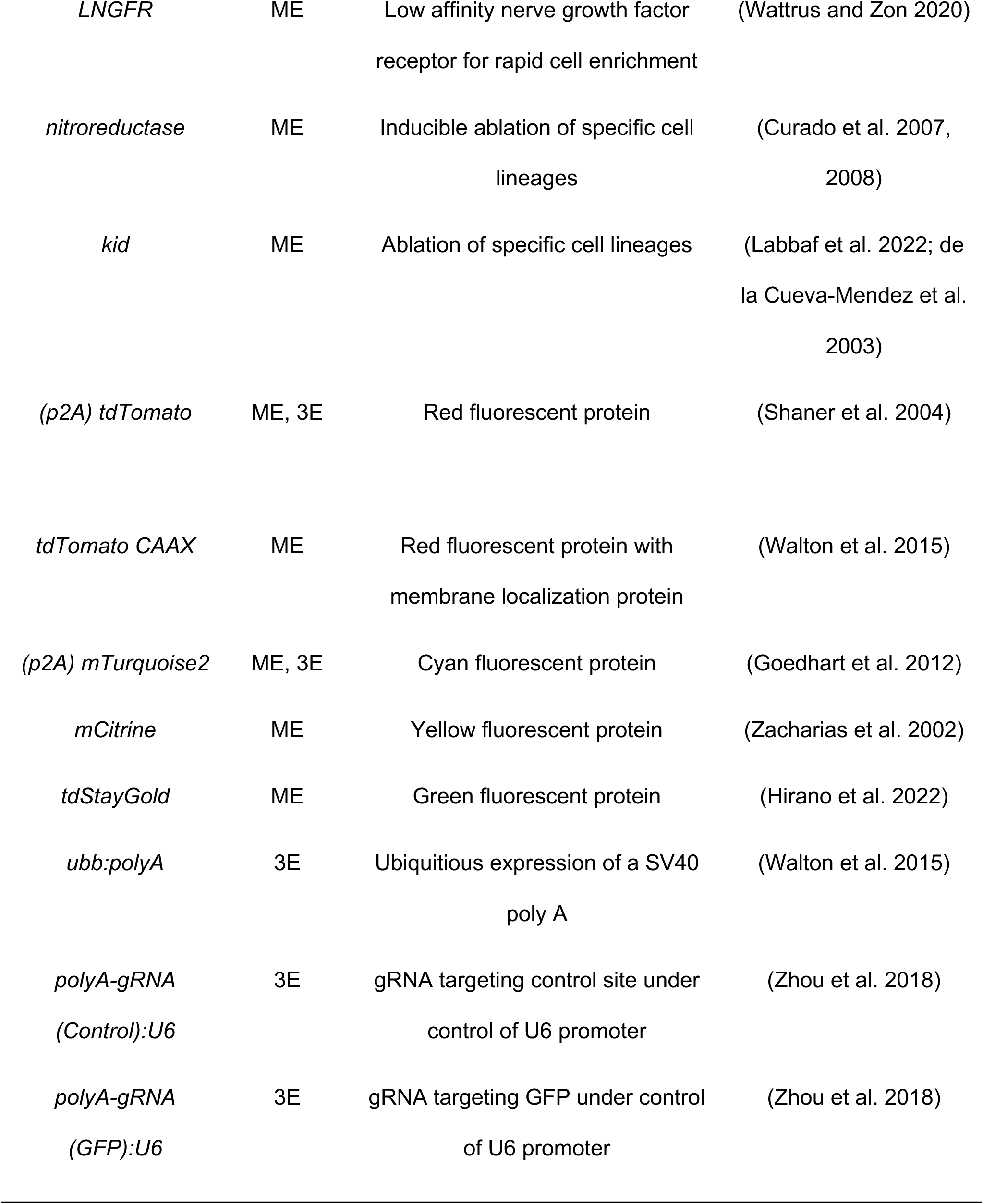
Library of insertion constructs.

| Insertion Elements | Position | Function | Reference |
| --- | --- | --- | --- |
| *irg1* | 5E | Infection-inducible macrophage-specific promoter | (C. J. Hall et al. 2013) |
| *mfap4* | 5E | Macrophage-specific promoter | (Walton et al. 2015) |
| *mpeg1.1* | 5E | Macrophage-specific promoter | (Ellett et al. 2011) |
| *lyz* | 5E | Neutrophil-specific promoter | (C. Hall et al. 2007) |
| *ubiquitin B (ubb)* | 5E | Promoter for ubiquitous expression | (Mosimann et al. 2011) |
| *ubb:loxP GFP loxP* | 5E | Promoter for ubiquitous expression of floxed GFP for use in combination with Cre | (Mosimann et al. 2011) |
| *hsp70* | 5E | heat shock protein 70-like promoter for inducible gene expression | (Shoji et al. 1998; Halloran et al. 2000; Kwan et al. 2007) |
| *lck* | 5E | T cell-specific promoter | (Langenau et al. 2004) |
| *runx1 + 23* | 5E | Promoter for expression in hematopoietic stem cells | (Tamplin et al. 2015) |
| *icre* | ME | Cre recombinase for inducible recombination of DNA between loxP sites | (Shimshek et al. 2002) |
| *(p2A) dLanYFP* | 3E | Yellow fluorescent protein | (Shaner et al. 2013) |
| *mNeonGreen* | ME, 3E | Green/ yellow fluorescent protein | (Shaner et al. 2013) |
| *LNGFR* | ME | Low affinity nerve growth factor receptor for rapid cell enrichment | (Wattrus and Zon 2020) |
| *nitroreductase* | ME | Inducible ablation of specific cell lineages | (Curado et al. 2007, 2008) |
| *kid* | ME | Ablation of specific cell lineages | (Labbaf et al. 2022; de la Cueva-Mendez et al. 2003) |
| *(p2A) tdTomato* | ME, 3E | Red fluorescent protein | (Shaner et al. 2004) |
| *tdTomato CAAX* | ME | Red fluorescent protein with membrane localization protein | (Walton et al. 2015) |
| *(p2A) mTurquoise2* | ME, 3E | Cyan fluorescent protein | (Goedhart et al. 2012) |
| *mCitrine* | ME | Yellow fluorescent protein | (Zacharias et al. 2002) |
| *tdStayGold* | ME | Green fluorescent protein | (Hirano et al. 2022) |
| *ubb:polyA* | 3E | Ubiquitous expression of a SV40 poly A | (Walton et al. 2015) |
| *polyA-gRNA (Control):U6* | 3E | gRNA targeting control site under control of U6 promoter | (Zhou et al. 2018) |
| *polyA-gRNA (GFP):U6* | 3E | gRNA targeting GFP under control of U6 promoter | (Zhou et al. 2018) |

### Generation of new transgenic zebrafish lines using ImPaqT

To test our PaqCI-based Golden Gate assembly toolkit as well as generate new immunological zebrafish lines for the community, we used ImPaqT to assemble a number of new transgenic lines. Using the neutrophil-specific promoter *lyz*, we generated a T*g(lyz:mTurquoise2)^ber2^* construct through Golden Gate assembly of 5E *lyz*, ME *mTurquoise2* and 3E *ubb:polyA* to create the *lyz:*m*Turquoise2* transgene construct (Fig. 2A-B). The construct was coinjected with *Tol2* transposase mRNA into single cell of fertilized eggs and at 2 dpi we observed strong blue fluorescence signal within likely neutrophil populations in the CHT (Fig. 2C). Positive animals were raised to adulthood and screening of F1 progeny of these animals revealed robust expression of mTurquoise2 within putative neutrophil populations throughout the animal (Fig. 2D). Crossing to the already established neutrophil reporter *Tg(lyz:EGFP)^nz117^*(C. Hall et al. 2007) zebrafish confirmed specific expression of our transgene within neutrophils (Fig. S2).

**Figure 2:**
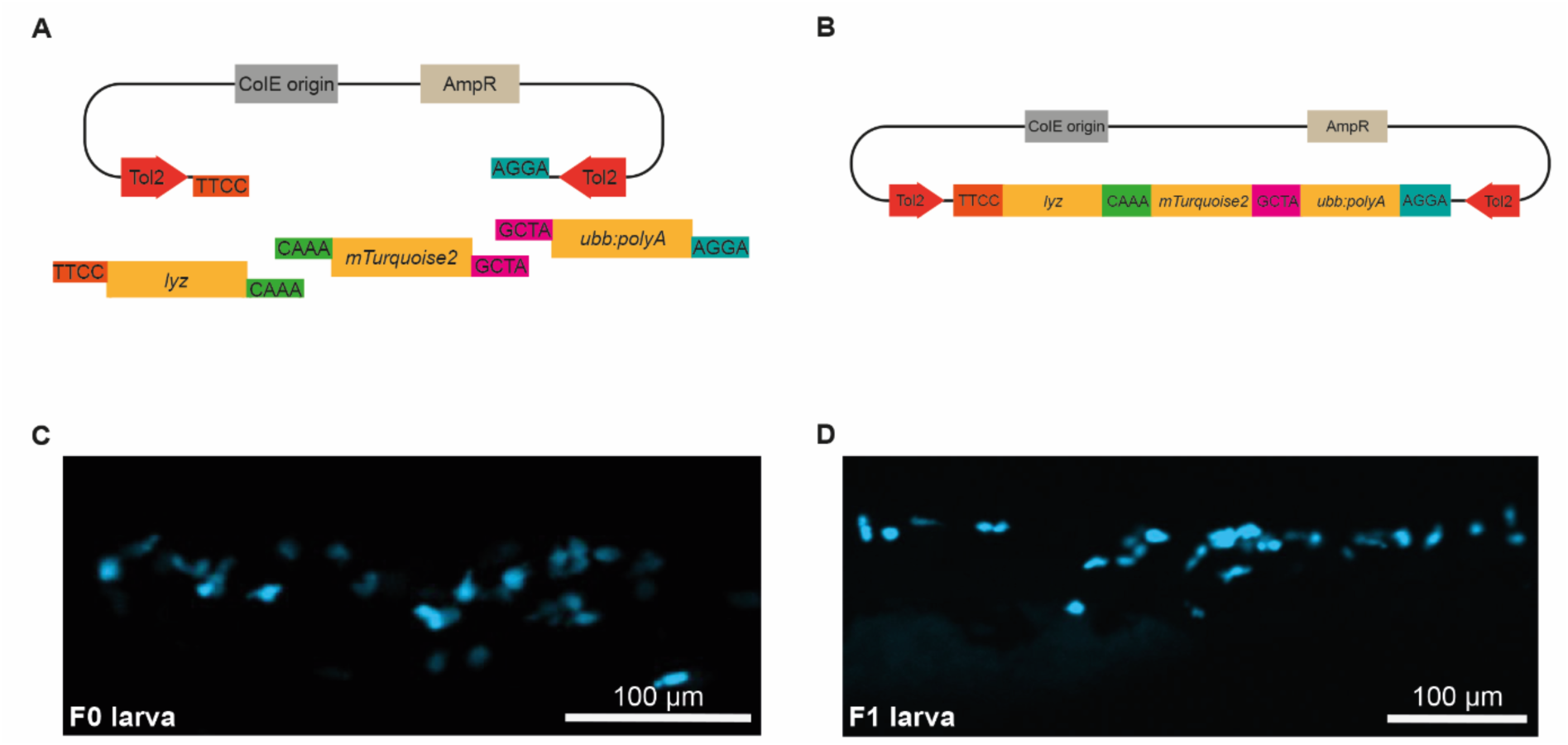
Generation of *Tg(lyz:mTurquoise2)* zebrafish A: Schematic of the Golden Gate assembly of *Tg(lyz:mTurquoise2)*. The backbone vector contains overhang sequences O1 (TTCC) and O4 (AGGA). *lyz* is a neutrophil-specific promoter containing overhang sequences O1 and O2 (CAAA). *mTurquoise2* encodes a bright cyan fluorescent protein with overhangs O2 and O3 (GCTA). *ubb:polyA* is assembled at the final position within the backbone with O3 and O4.B: Schematic of the final assembly product of *Tg(lyz:mTurquoise2)*. C-D: Images of a *Tg(lyz:mTurquoise2)* F0 (C) and F1 (D) larva showing neutrophil response in cyan at 2 days post fertilization.

Similarly, we used the ImPaqT toolkit to generate a new macrophage-specific transgene using the infection-inducible macrophage-specific promoter *irg1* and the recently described bright green fluorescent protein *tdStayGold* (Hirano et al. 2022). To generate the *Tg(irg1:tdStayGold)^ber3^* construct, the 5E *irg1,* ME *tdStayGold* and 3E *ubb:polyA* elements were combined into the backbone vector (Fig. 3 A-B). We injected zebrafish with this construct and as *irg1* is an inflammation responsive promoter that is only weakly expressed in the absence of immune stimulation, we injected larvae with LPS before screening. After LPS injection, we were able to identify a number of F0 founders with strong green fluorescence within subpopulations of their macrophages (Fig. 3C). These positive F0 larvae were raised to adulthood and screening of F1 progeny revealed expression of tdStayGold throughout macrophage populations upon stimulation with LPS (Fig. 3 D).

**Figure 3:**
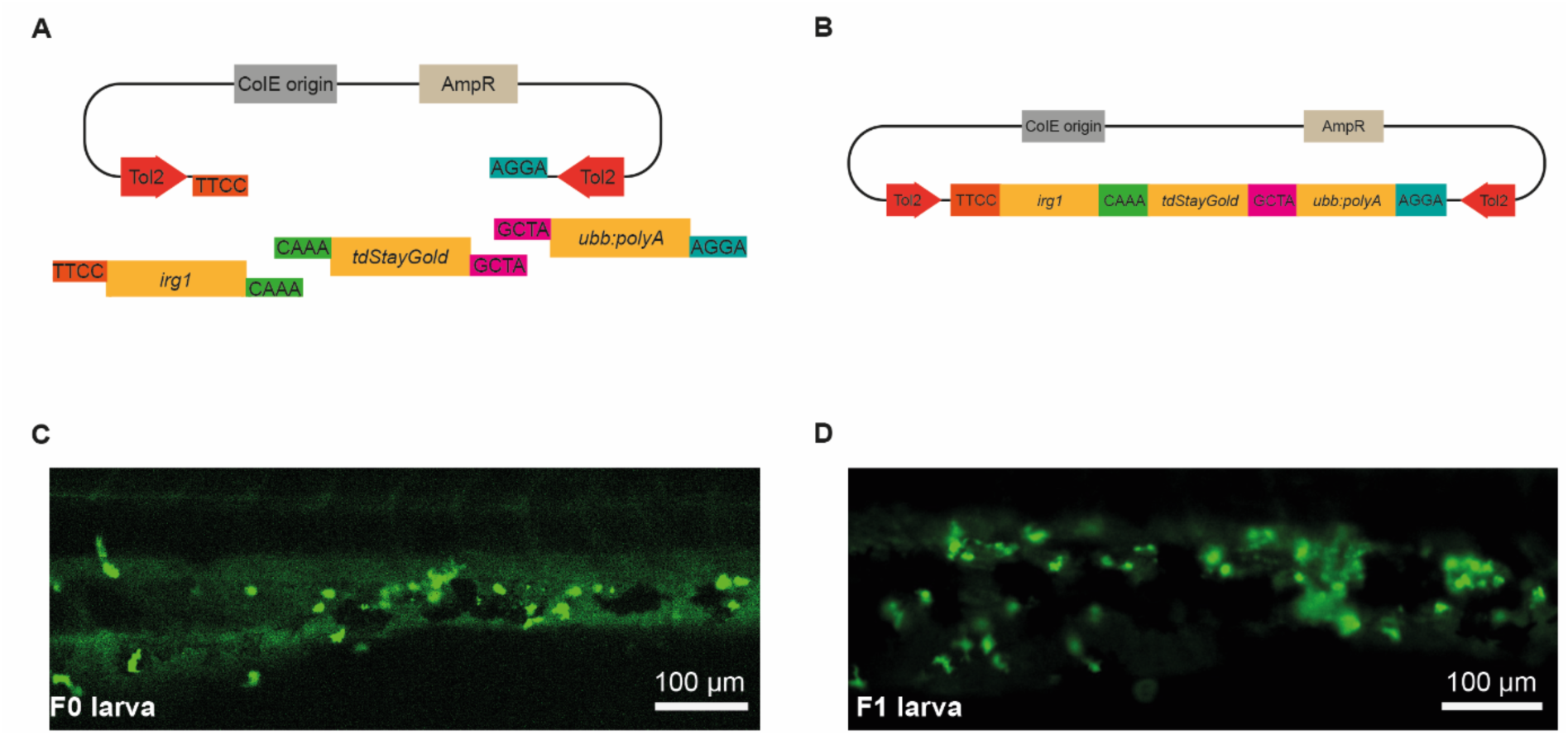
Generation of *Tg(irg1:tdStayGold)* zebrafish A: Schematic of the Golden Gate assembly of *Tg(irg1:tdStayGold)*. The backbone vector contains overhang sequences O1 (TTCC) and O4 (AGGA). *irg1* is a macrophage-specific promoter containing overhang sequences O1 and O2 (CAAA). *tdStayGold* encodes a bright green fluorescent protein with overhangs O2 and O3 (GCTA). *ubb:polyA* is assembled at the final position within the backbone with O3 and O4. B: Illustration of the final assembly product of *Tg(irg1:tdStayGold)*. C-D: Images of a *Tg(irg1:tdStayGold)* F0 (C) and F1 (D) larva showing macrophage-specific tdStayGold expression.

We also aimed to test the utility of the tools that we have designed within our kit. To these ends, we created an infection-inducible, macrophage specific *icre* transgene construct, *Tg(irg1:icre-p2A mTurquoise2)*. This construct was generated by combining the 5E *irg1,* ME *icre* and 3E *p2A mTurquoise2* (Fig. 4A and 4B). Animals injected with this transgene showed macrophage-specific mTurquoise2 signal after injection with LPS (Fig. 4C). To further confirm that this construct was able to drive rearrangement of floxed reporters, we injected *Tg(-3.5ubb:LOXP-EGFP-LOXP-mCherry)^cz1701^* (Mosimann et al. 2011) zebrafish with this construct. These animals were injected with LPS and imaged for 48 h to visualize rearrangement of the floxed construct. LPS treatment drove the induction of *icre* and the co-translationally cleaved *p2a mTurquoise2* transgene within macrophages starting ca. 4 h post LPS injection. While the induction of the *irg1:icre-p2a mTurquoise2* transgene was transitory, this short-term induction of Cre could induce permanent rearrangement of the *loxP-EGFP-loxP-mCherry* construct as seen by induction of red fluorescent mCherry within macrophage populations (Fig. 4D, S3, Movie S1). Control animals without the *icre*-construct neither showed mTurquoise2 nor mCherry signal in macrophages (Fig. S3, Movie S 2). This experiment showed not only the functionality of the individual insertion constructs but also the range of assembly constructs that can be generated with them.

**Figure 4:**
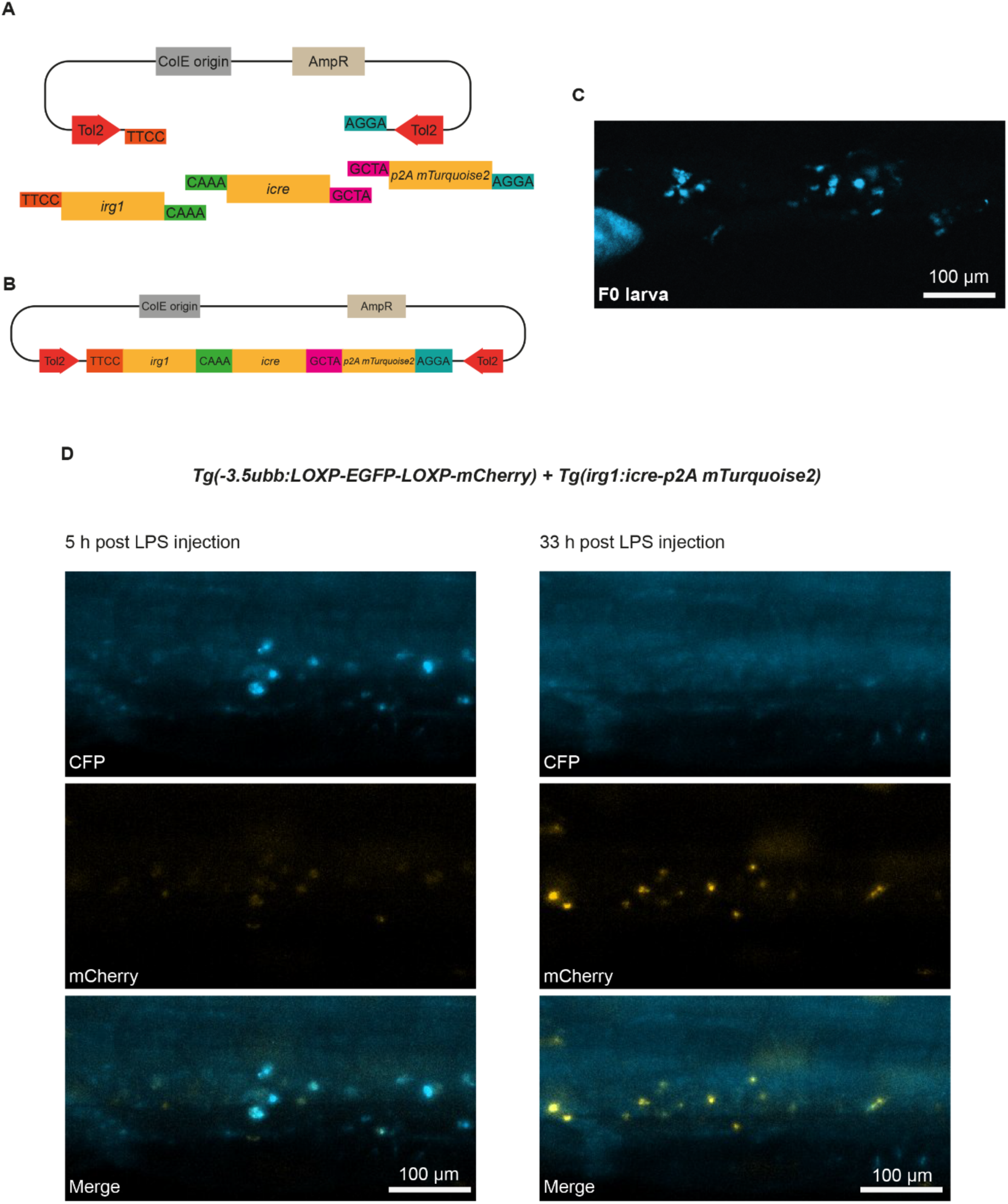
Generation of *Tg(irg1:icre-p2A mTurquoise2)* zebrafish for lineage tracing of individual macrophages A: Schematic of the Golden Gate assembly of *Tg(irg1:icre-p2A mTurquoise2)*. The backbone vector contains overhang sequences O1 (TTCC) and O4 (AGGA). The macrophage promoter *irg1* contains overhang sequences O1 and O2 (CAAA). *icre* contains overhangs O2 and O3 (GCTA) for the middle position. *p2A mTurquoise2* is assembled at the final position within the backbone with O3 and O4. B: Illustration of the final assembly product of *Tg(irg1:icre-p2A mTurquoise2)*. C: Image of a *Tg(irg1:icre-p2A mTurquoise2)* F0 larva showing macrophage-specific mTurquoise2 signal after stimulation with LPS. D: Images of a *Tg(-3.5ubb:LOXP-EGFP-LOXP-mCherry)* larva that was introduced to the *Tg(irg1:icre-p2A mTurquoise2)* construct at single cell stage and was treated with PTU and LPS. The images show a macrophage-specific mTurquoise2 signal (e.g. 5 h post LPS injection) and a macrophage-specific mCherry signal (e.g. 33 h post LPS injection).

### ImPaqT is expandable through the introduction of new overhang sequences

A limitation of many existing systems for transgenesis is that these systems have a fixed number of elements that can be combined to generate transgenes. Golden Gate methods however, are expandable through the addition of new overhangs and have been used to join 10-24 or even more than 50 fragments of DNA in a single reaction (Kucera 2021; Pryor et al. 2022). The ability to expand this kit to include larger numbers of constructs would simplify the generation of complex transgenes including multifragment promoters, loxP sites, localization tags (e.g NLS or prenylation signal) or multigene constructs for example. To provide proof of concept for the expandability of ImPaqT, vectors were designed to enable the construction of a *ubb* promoter-driven construct from three similar sized *ubb* promoter fragments, *ubb_1_, ubb_2_* and *ubb_3_*. These fragments were constructed such that the first fragment (*ubb_1_*) and the last fragment (*ubb_3_*) use the same O1 and O2 overhangs used by 5E constructs so that the array of promoter fragments can be cloned into the existing position of 5E elements and used alongside already existing ME and 3E constructs (Fig. 5A-B). New internal overhangs were designed for the *ubb_1_, ubb_2_* and *ubb_3_* fragments to combine these distinct fragments (Fig. 5A-D). Individual *ubb* fragments were used for Golden Gate assembly in combination with ME *mTurquoise2* and 3E *ubb:polyA* (Fig. 5C) which led to the expected transgenesis construct *Tg(ubb:mTurquoise2)^ber4^*(Fig. 5D) that was injected into embryos at single cell stage. Screening of injected animals confirmed that transgenic embryos established with this construct had the expected expression pattern (Fig. 5E). We also found robust expression in the F1 progeny (Fig. 5F) after raising positive F0 animals, confirming that injected animals are viable and able to pass on the transgene to the next generation. Together, these findings demonstrated proof-of-concept for the expansion of this transgene assembly system enabling users to add fragments as needed through the addition of novel unique 4-base overhangs while maintaining compatibility with existing ImPaqT constructs.

**Figure 5:**
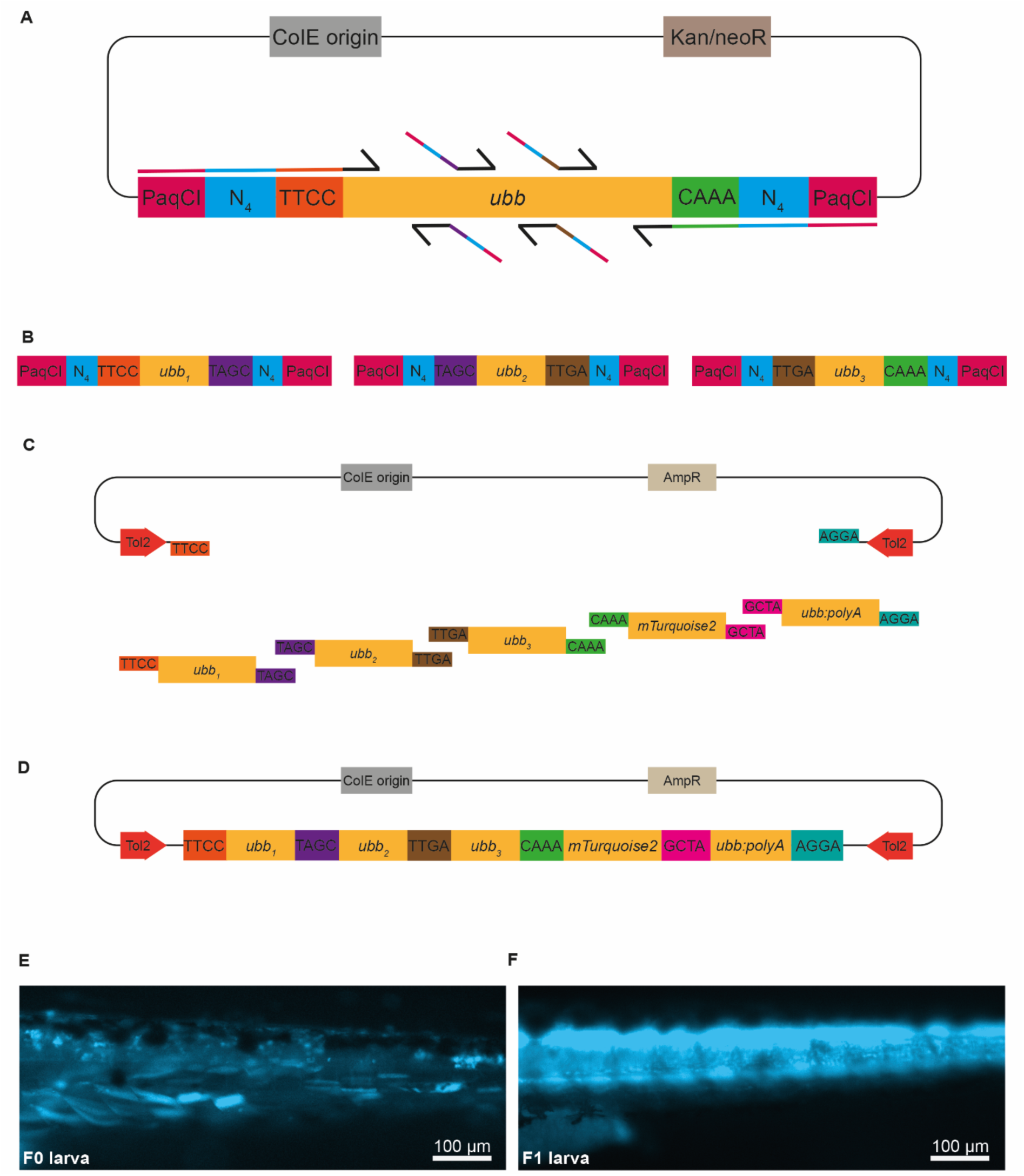
**ImPaqT allows for expandability of individual positions while retaining compatibility with already established transposon components.** A: Schematic of the 5E *ubiquitin B* (*ubb*) promoter construct containing PaqCI recognition sites (PaqCI) along with 4-base spacer sequences (N_4_) and two overhangs (O1: TTCC, O2: CAAA) for positioning at the 5’ end of the assembly construct. Primers were designed to break up *ubb* into three fragments with new overhang sequences. B: Schematic of PCR amplified *ubb* fragments *ubb_1-3_* with PaqCI, N_4_ and O1 and O1A (TAGC), O1A and O1B (TTGA), and O1B and O2, respectively. C: Schematic of the Golden Gate assembly of 5E *ubb_1-3_*, ME *mTurquoise2* and 3E *ubb:polyA* into the backbone vector. D: Illustration of the final assembly product of *Tg(ubb_1-3_:mTurquoise2)*. E-F: Images of a *Tg(ubb:mTurquoise2)* F0 (E) and F1 (F) larva showing ubiquitous mTurquoise2 signal.

### ImPaqT allows for rapid transgene generation through cloning from PCR

Within the framework of ImPaqT, insertion elements are PCR amplified for the addition of PaqCI sites, spacer sequences and overhangs that are required for the Golden Gate assembly reaction, and subsequently subcloned into vectors prior to the assembly of the transgene. This subcloning step not only ensures an error-free and stable storage of these constructs but also allows for the generation of a stable library of elements that can be easily combined, re-used and shared within the community. However, the additional cloning step is also more time consuming and, in rare cases where the construct is toxic to bacteria, can result in alteration of parts of the construct. In individual experimental strategies, e.g. for testing new elements or for the single use of individual elements, it may be redundant to subclone and store the PCR product. Instead, the Golden Gate assembly can be performed directly with PCR products, resulting in faster and more flexible transgene generation. To demonstrate, that ImPaqT also allows for rapid creation of functional transgenes by cloning directly from PCR, we assembled *Tg(mpeg1.1:tdTomato-p2A-rac2^WT^)* and *Tg(mpeg1.1:tdTomato-p2A-rac2^D57N^)* by using PCR products directly for the Golden Gate assembly reaction without subcloning into vectors. Rac2 is a small GTPase involved in immune cell migration (Salas-Vidal et al. 2005) and dominant-negative mutations of Rac2 including the patient-derived D57N mutation are linked with migration arrest in immune cells (Deng et al. 2011). Here, we used the macrophage-specific *mpeg1.1* promoter to drive tdTomato followed by a self-cleaving peptide and then either Rac2 (Rac2^WT^) or a dominant-negative mutant form of Rac2, Rac2^D57N^. N-terminal tagging of Rac2 is necessary as Rac2 undergoes a C-terminal prenylation event which is required for its membrane localization and activity. As this approach required us to invert our usual N-terminal gene, C-terminal fluorescent protein order, the flexibility afforded by PCR cloning simplified the construction of this swapped gene order. To generate these two transgenes, PCR products encoding the macrophage-specific promoter *mpeg1.1* with overhangs O1 and O2, *tdTomato* with overhang O2 and O3, and p2A with overhang O3 and O3B (CTCG) were combined together with a dsDNA fragment (IDT gBlock) encoding either wildtype or dominant-negative *rac2* with overhangs O3B and O4 (Fig. 6A, S5D-E). These fragments were cloned together with the already existing backbone vector to generate *Tg(mpeg1.1:tdTomato-p2A-rac2^WT^)* and *Tg(mpeg1.1:tdTomato-p2A-rac2^D57N^)* (Fig. 6A-C). The final transgene construct was injected into embryos at single cell stage. To observe macrophage migration, at 2 dpf embryos were anesthetized and the tip of the tail fin was cut with a scalpel to create a wound. Subsequent imaging demonstrated that while in *Tg(mpeg1.1:tdTomato-p2A-rac2^WT^)* embryos macrophages readily migrated to the wound in the tail fin 2 h post wounding (Fig. 6D), in *Tg(mpeg1.1:tdTomato-p2A-rac2^D57N^)* animals, macrophages were frequently observed throughout the body but were not seen migrating to the tailwound throughout imaging (Fig. 6E). Together, this data provided proof of concept that ImPaqT can be used for rapid generation of functional transgenes and also offered new insights about the utility of this toolkit for immunological research.

**Figure 6:**
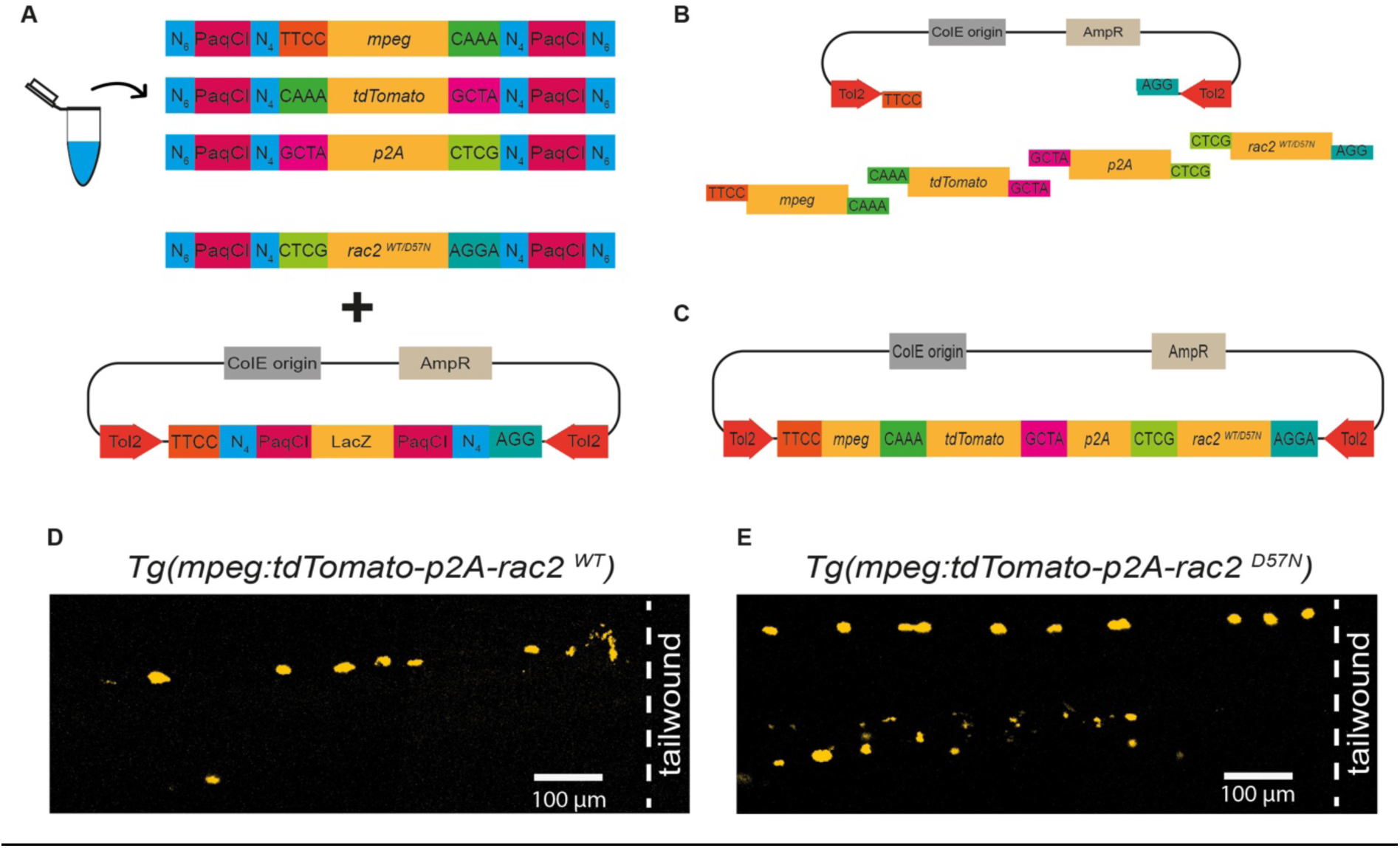
**ImPaqT allows for rapid transgene generation by Golden Gate assembly from PCR.** A: Schematic of PCR amplified fragments *mpeg1.1*, *tdTomato*, and *p2A* containing PaqCI, N_4_ and overhangs O1 (TTCC), O2 (CAAA), O3 (GCTA) and O3B (CTCG) and dsDNA fragment encoding *rac2^WT/D57N^* containing O3B and O4 (AGGA). B: Schematic of the Golden Gate assembly of *mpeg1.1, tdTomato, p2A and rac2^WT/D57N^* into the backbone vector. C: Illustration of the final assembly products *Tg(mpeg1.1.tdTomato-p2A-rac2^WT/D57N^)*. D-E: Images of a *Tg(mpeg1.1.tdTomato-p2A-rac2^WT^)* F0 (D) and *Tg(mpeg1.1.tdTomato-p2A-rac2^D57N^)* F0 (E) larva 2 h post tail wounding showing macrophages.

## Discussion

Transgenic animals remain essential for many research questions. They enable the visualization of cells and tissues, molecular pathways, immune responses and the perturbation of these pathways to identify the molecular mechanisms driving pathology. Use of the zebrafish as a model organism has been steadily increasing in many research disciplines due to the unique advantages of this model. This is true in immunology and host-pathogen interactions, where the optical transparency afforded by the zebrafish model have led to many research discoveries, but where, as a new immunology model, there are limited transgenic tools. Here, we demonstrate a new cloning toolkit for the generation of immunological transgenes that is based on Golden Gate cloning. The Golden Gate approach results in shorter overlap regions (4 bp) than recombination-based techniques like Gateway cloning, while compared to seamless, homology based cloning methods such as Gibson cloning, it allows for straightforward modular cloning approaches. The Golden Gate approach also takes advantage of off-the-shelf restriction enzymes and ligase, compared to other cloning approaches which use proprietary enzymes for cloning.

We have termed this toolkit ImPaqT - **Im**munological toolkit for **Paq**CI-based Golden Gate Assembly of Tol2 **T**ransgenes. It includes a range of immunologically focused promoters along with genetic tools and fluorescent proteins that can be assembled using PaqCI-based Golden Gate cloning. These tools will enable the flexible assembly of transgenes to drive expression and to genetically manipulate, purify and ablate immune cell populations of interest. While zebrafish possess the same immune lineages of other vertebrates, as a relatively new model for immunology studies, the zebrafish has a more limited array of cell type-specific promoters than mouse models. For instance, despite the identification of potential NK- and ILC-specific (Hernández et al. 2018) as well as mast cell-specific (Dobson et al. 2008) genes, promoters specific for these cell populations have not been described. As new immune-specific promoters are identified, these can be readily added to this toolkit. While we have targeted the immune system as a particular area of need, this Golden Gate approach can also be adapted to any other tissue or cell lineages within the zebrafish.

While Golden Gate cloning enables the use of shorter homology stretches and the combinatorial use of constructs for the assembly of diverse transgenes, as a restriction enzyme-based system, a limitation of Golden Gate systems, compared to other assembly approaches, has been the risk of off-target cutting within the cloned sequences. This off-target cutting requires mutation of insert sequences to eliminate these sites (termed domestication) (Marillonnet and Grützner 2020; Bird et al. 2022; Grützner and Marillonnet 2020), which can potentially alter function of domesticated promoters and genes. While domestication can result in changes in the sequence that may alter behavior of the genetic element, these in principle can be managed for genes of interest using silent mutations in critical bases. Commonly, endogenous PaqCI sites can be mutated by PCR amplification of the element using primers designed to incorporate a silent mutation in the PaqCI site (e.g. CACCT**<u>G</u>**C is changed to CACCT**<u>C</u>**C) (Fig. S6A), followed by In-Fusion cloning to reconstitute the construct (Fig. S6B-C). Alternatively, domestication can be performed in other ways, for example by using overlap extension PCR or QuickChange. While promoters cannot be managed through the same approach, promoter fragments used for zebrafish are frequently large (commonly 2-6 kb) and it is unlikely that the mutation of a single base pair will inadvertently ablate the promoter activity. Consistent with this, we have not seen any changes in promoter activity or localization in the two promoters that we domesticated for the ImPaqT toolkit, although further cloning of additional promoters will more completely test this idea. Our choice of PaqCI itself seeks to address the limitations of domestication, as PaqCI uses a 7 base recognition site. In contrast to the 6 base-cutters that have been described in previous zebrafish Golden Gate approaches (Kirchmaier et al. 2013; Jiang et al. 2022), the rare cutting of PaqCI (if random, 1 in 16,384 bases) makes this enzyme better suited for the large DNA fragments that are frequently used for zebrafish promoters.

As a Golden Gate-based toolkit, ImPaqT introduces only short (4-base) overlap regions between individual elements, in contrast to recombination-based systems like Gateway that require the addition of long (20-25 bp) sequences. However, it must be considered that the addition of 4 extra bases could still potentially lead to side effects on the endogenous functions of promoters or proteins. While it is unlikely that the addition of 4 bases for promoters or 2 codons for proteins has detrimental effects, for some proteins that require free C- or N-termini, these methods could be disruptive and other methods of targeting, including internal tagging or transgenesis markers, need to be used. In critical cases, overhang sequences can be designed to match the sequence of the final product, ensuring that there is ultimately no sequence change from the endogenous sequence (Bird et al. 2022) at the cost of reduced flexibility. Similarly in the case of promoters, the 4 bp addition could rarely result in the creation of novel transcription factor binding sites, given the short sequences that are recognized by these proteins. This might be magnified at the transcriptional start site, given the density of regulatory elements here. Importantly, consistent with the relative rarity of these risks to both proteins and promoters, we have not observed any side effects on endogenous functions of promoters or proteins in our study.

A limitation of existing systems has been the use of a fixed number of constructs for assembly of the transgene. One benefit of the Golden Gate directed system is that it can be expanded for cloning of complex transgenes, with some Golden Gate assemblies encompassing 20 or even more than 50 fragments (Kucera 2021; Pryor et al. 2022). While we have designed the system to incorporate 4 distinct DNA segments (3 inserts and a backbone vector) the flexibility of Golden Gate cloning means that our system can be readily expanded and contracted while maintaining compatibility with the other components of the kit. As proof-of-concept, we were able to break a single 5E encoding the *ubb* promoter into 3 distinct plasmids that we could subsequently assemble together with existing ME, 3E and backbone elements to generate a functional *ubb* driven transgene. The flexibility of this system will simplify the construction of large and complex transgenes. Potential applications would include breaking up larger promoter fragments into smaller and easier to clone elements, insertion of recombination sites for enzymes like Cre or Flp for rearrangement, assembly of multipromoter constructs encoding genes such as Cas9 and CRISPR gRNAs for cell type-specific knockout, or the inclusion of epitope or localization tags. Our largest reaction comprised 6 fragments (Fig. 5), although in theory, ImPaqT allows for the assembly of even more elements than investigated here. While larger constructs are likely possible, in our experience 6 fragments is a sufficient level of complexity for most transgenes.

In addition, we have also showed, that ImPaqT can be used to rapidly generate functional transgenes by Golden Gate assembly of PCR products and larger dsDNA fragments that can be readily ordered from suppliers without the need to subclone insertion elements into vectors prior to the assembly. The Golden Gate assembly of PCR products saves time and can be useful for rapid testing of new elements or for the single use of individual constructs, when long-term storage, multiple uses or sharing is not planned.

Lastly, while we have applied this Golden Gate method for the construction of Tol2 transgenes, the advantages of the PaqCI-Golden Gate method will likely be useful for the editing of zebrafish by other approaches as well. For instance, CRISPR insertion has been increasingly used to generate knock-in zebrafish lines, via either homologous recombination, non-homologous methods or by using short homology stretches. Knock-in approaches require the construction of donor vectors which incorporate varied sequences such as homology regions, fluorescent proteins and stretches of genomic DNA (Albadri et al. 2017). For the application of ImPaqT for CRISPR/Cas9-mediated gene editing, we also incorporated the element polyA-gRNA:U6 with a gRNA targeting either a control site (Zhou et al. 2018) or GFP that can be used as either a non-targeting gRNA or to assess the efficiency of gene disruption in GFP-expressing populations with both under the control of the U6 promoter. The construct is included in the library as a 3’ element but with overhang sequences CGTA (O3C) and O4 to enable the more complex constructions that are frequently seen in these constructs (e.g. a tissue-specific promoter, cas9, a fluorescent protein and the U6-promoter-gRNA cassette). The gRNA sequences of 3E polyA-gRNA:U6 can easily be exchanged by homology-directed cloning to address new targets. Similarly, many groups also use I-SceI insertion approaches to generate transgenic zebrafish (Hoshijima et al. 2016). The PaqCI Golden Gate method could be adapted to simplify the construction of transgenes and editing constructs for these other approaches as well. Although the Golden Gate approach is similarly efficient to other methods to generate transgenesis constructs, the minimal additional sequence, and the ability of the Golden Gate method to expand and contract as needed provide flexibility that is beneficial for all of these editing and transgenesis methods. Our efforts here have been focused on immune populations, but the toolkit can also be broadened for the use beyond the immune system. In fact, with the addition of new cell-type specific promoters, the tools in ImPaqT can be used in other fields, such as developmental biology, regeneration and neuroscience, in which the zebrafish has been a popular model organism.

## Materials and Methods Ethics statement

Zebrafish husbandry and experiments were approved by the “Landesamt für Gesundheit und Soziales” (LaGeSo) Berlin (Germany).

## Zebrafish husbandry and handling

Zebrafish (*Danio rerio*) embryos were raised in petri dishes containing E3 medium at 28°C for up to five days. Animals older than 5 days were maintained in a circulating system with constant water conditions of 28°C, pH 7.0 - 7.3, and a conductivity of 600-700 µS on a 12-hour light/ 12-hour dark cycle. They were fed twice a day with artemia and once a day with dry food.

## Generation of transgenic zebrafish lines

### Zebrafish strains

All zebrafish strains, including *Tg(lyz:mTurquoise2)^ber2^, Tg(irg1:tdStayGold)^ber3^*, and *Tg(ubb:mTurquoise2)^ber4^*were in the *AB wildtype background and generated in this study by applying the Tol2 transgenesis system as previously described (Kawakami et al. 2004).

Transient injection of *Tg(irg1:icre-p2A-mTurquoise2)* and *Tg(mpeg1.1:tdTomato-p2A-rac2^WT/D57N^)* were also done in *AB background. *Tg(-3.5ubb:LOXP-EGFP-LOXP-mCherry)^cz1701^* zebrafish were obtained from the European Zebrafish Resource Center (EZRC). *Tg(-3.5ubb:LOXP-EGFP-LOXP-mCherry)^cz1701^*(Mosimann et al. 2011) and *Tg(lyz:EGFP)^nz117^* (C. Hall et al. 2007) have been previously described.

### Preparation of the backbone vector and inserts

The backbone vector containing the LacZ gene flanked by PaqCI sites and *Tol2* arms in a vector backbone with a pMB1 origin and an AmpR gene for bacterial selection (pTWIST Amp High Copy) was purchased from Twist Bioscience. The complete sequence is shown in Figure S4. Primer sequences are shown in Table S1. Gene fragments for *runx1+23* (with the endogenous PaqCI site mutated), *tdStayGold* and truncated human *LNGFR* (zebrafish codon optimized and with endogenous PaqCI sites mutated) were purchased from Twist Bioscience and TOPO cloned into pCRII TOPO (Thermofisher Zero Blunt TOPO). The sequence for *rac2^WT^* and *rac2^D57N^* containing PaqCI sites, spacer sequence (N_4_), overhangs O3B and O4 as well as 6-8 additional bases at the 5’ and 3’ ends (N_6-8_) and the sequence for the element polyA-gRNA(Control):U6 and polyA-gRNA(GFP):U6 (Addgene Plasmid #107599) containing PaqCI sites, N_4_, overhangs O3C and O4 were purchased as gBlocks from IDT. The complete sequences are shown in Figure S5.

PaqCI recognition sites with 4-base spacer sequences and 4-base overhangs were added to DNA sequences via PCR amplification. Primer were designed to add PaqCI sites such as cleavage resulted in a gene fragment with a 4-base overhang that is compatible to the overhang of another gene fragment that will be adjacent to it in the final assembly. Appropriate 4-base overhang sequences were confirmed using the online tool NEBridge Ligase Fidelity Viewer^™^. PCR amplification was performed using Q5 polymerase and analyzed by gel electrophoresis. The PCR product was cleaned up or, in case of multiple bands, the band of correct size was excised under 70% UV light using a clean scalpel and DNA was extracted using Nucleo Spin Gel and PCR Clean-up Kit (Macherey-Nagel). DNA sequences were confirmed by sequencing. Gel purified gene fragments were subcloned into a Zero Blunt TOPO vector (Invitrogen) and transformed into competent NEB 5α *E. coli* cells (C2987H). Cells were cultivated in LB medium containing 50 µg/ml Kanamycin and plasmids were isolated using the Nucleo Spin Plasmid Quick Pure protocol (Macherey-Nagel). For the Golden Gate assembly of transgenes from PCR, inserts were PCR amplified with an additional 6-8 bases added to the 5’ and 3’ end for efficient restriction digestion and Golden Gate assembly directly from PCR.

Both the *ubb* and *lck* promoters had endogenous PaqCI cleavage sites in the promoter sequence. To domesticate these promoters, we mutated these PaqCI sites with the indicated PCR primers (Table S1) and assembled them with In-Fusion cloning (Takara Bio).

### Golden Gate based assembly of expression constructs

Golden Gate assembly was performed according to the NEB protocol. In detail, 75 ng of the backbone vector and each insertion construct was mixed with 10 U PaqCI, 0.5 µl PaqCI activator (20 µM) and 400 U T4 DNA ligase in T4 DNA ligase buffer to a final volume of 20 µl. Final assembly product was obtained after 60 Cycles of incubation at 37°C for 1 min and incubation at 16°C for 1 min, followed by 5 min incubation at 37°C and 5 min incubation at 60°C using a thermocycler. The final product was transformed into competent NEB 5α *E. coli* cells (C2987H) and cells were cultivated in LB medium containing 20 µg/ml Ticarcillin. Plasmids were isolated using the Nucleo Spin Plasmid Quick Pure protocol (Macherey-Nagel) and PCR purified using the Nucleo Spin Gel and PCR Clean-up Kit (Macherey-Nagel) before microinjection.

### Microinjections

The injection mixture contained 25 ng/µL polyadenylated mRNA encoding the Tol2 transposase, 50 ng/µL transposon donor plasmid and 10% phenol red in a final volume of 10 µl 1 X Tango Buffer (Thermo Fisher). The mMessage mMachine kit (Life Technologies) was used to transcribe the mRNA from T3TS-Tol2 (Balciunas et al. 2006). A volume of ∼1 nL was injected into the single cell of fertilized eggs by using a microinjector.

### Imaging

Injected larvae were screened for insertion of the transgene 2 days post injection (dpi) using a Nikon ECLIPSE Ti2 microscope (Nikon) with a Dragonfly 200 confocal system (Oxford Instruments). Animals that were positive for the transgene were raised to adulthood and outcrossed to the *AB wildtype strain for selection of transgenic founder zebrafish. Before screening *Tg(irg1:icre-p2A mTurquoise2)* and *Tg(irg1:tdStayGold)* embryos, the activity of the *irg1* promoter was induced by injection of LPS. For that purpose, animals were anesthetized with tricaine (MS-222) at a final concentration of 0.016% and injected into the caudal vein with LPS (Sigma-Aldrich, F3665) (final: 0.8 mg/ml). Screening was performed at least 4 h after injection. To reduce pigmentation, *Tg(-3.5ubb:LOXP-EGFP-LOXP-mCherry)* embryos were treated with 45 µg/ml 1-phenyl-2-thiourea (PTU) starting ca. 20 h post fertilization. To visualize macrophage migration in *Tg(mpeg:tdTomato-p2A-rac2^WT/D57N^)* embryos, animals at 2 dpf were anesthetized with 0.016% tricaine and a piece of the tail fin was cut with a scalpel, followed by imaging 2 h post transection. Image analysis was performed using the Fiji ImageJ software (National Institute of Health, USA).

### Line and reagent availability

All new lines and plasmids are available from the corresponding author upon contact and will also be deposited to AddGene.

## Acknowledgements

We would like to thank Miriam Herbert for sharing the ME *kid* construct. Additionally, we would like to thank our animal welfare officer Antje Dreyer and our animal caretakers Silke Lehmann, Marten Walk, Carsten Weiland, Ines Neumann, Jens-Phillip Otto und Janine Bleske for their great work and support.

## Data availability

All relevant data can be found within the article and its supplementary information.

