## Supplementary Figures and Table for "ImPaqT - A Golden Gate-based Toolkit for Zebrafish Transgenesis"

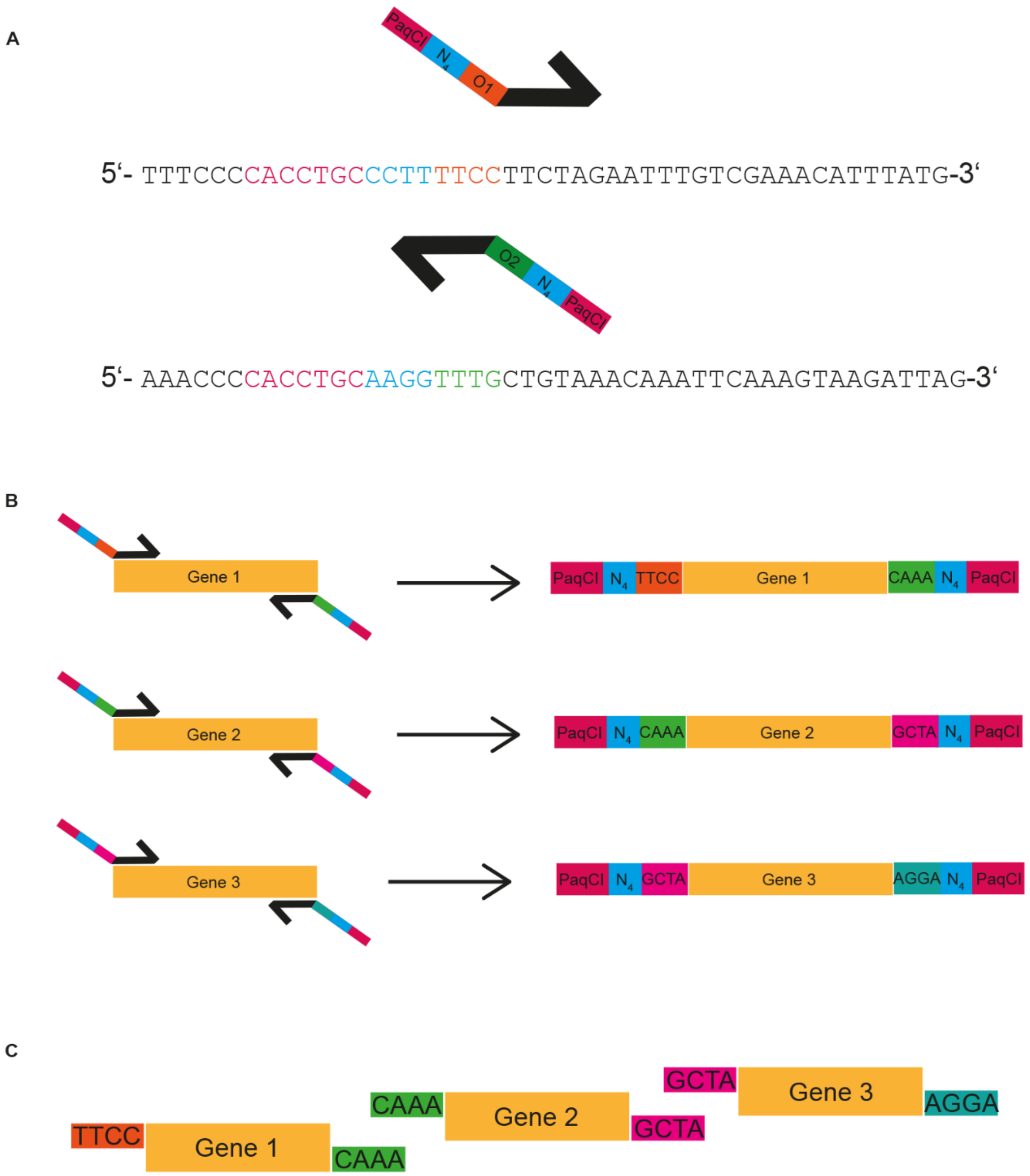


**Supplementary Figure 1: Preparation of insertion constructs**

A: Schematic of primer design for generating a 5E insertion containing overhang sequences O1 (TTCC, orange) and O2 (TTTG, green), 4-base spacer sequence ($N_4$, CCTT/AAGG, blue) and PaqCI recognition site (CACCTGC, pink) with example primer sequence for the 5E *ubiquitin B* promoter insertion. B: Schematic of the amplification of genes for generation of 5E, ME ad 3E with primers containing the PaqCI recognition site, $N_4$ and a specific overhang sequence. C: Schematic of digested insertion elements with specific overhangs.

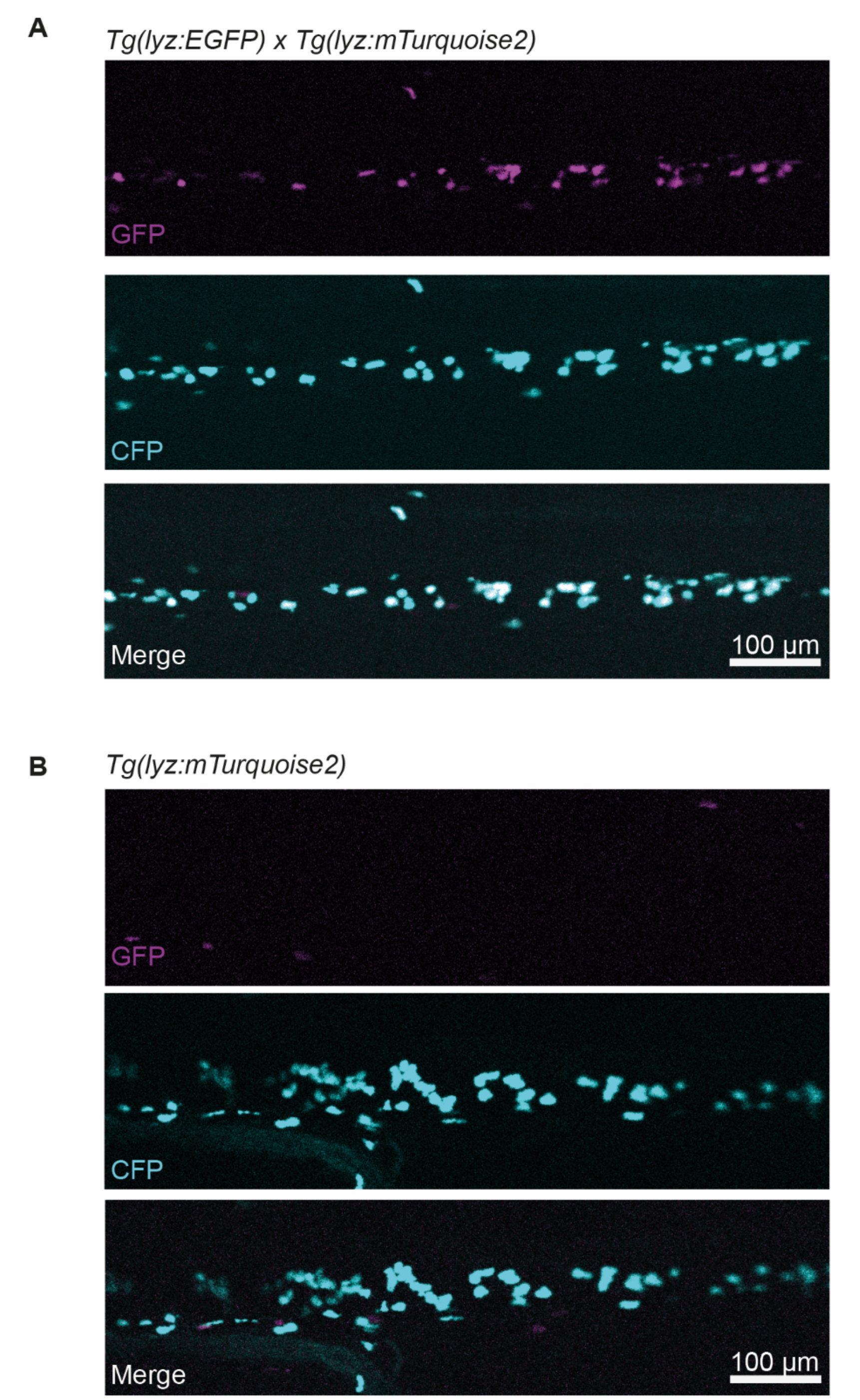


**Supplementary Figure 2: Verification of the *Tg(lyz:mTurquoise2)* zebrafish line**
*Tg(lyz:EGFP)* and *Tg(lyz:mTurquoise2)* zebrafish were crossed. Images of a double-positive larva are shown in A and images of a *Tg(lyz:mTurquoise2)* single-positive larva in B. GFP is presented in magenta, CFP signal in cyan and merge in white.

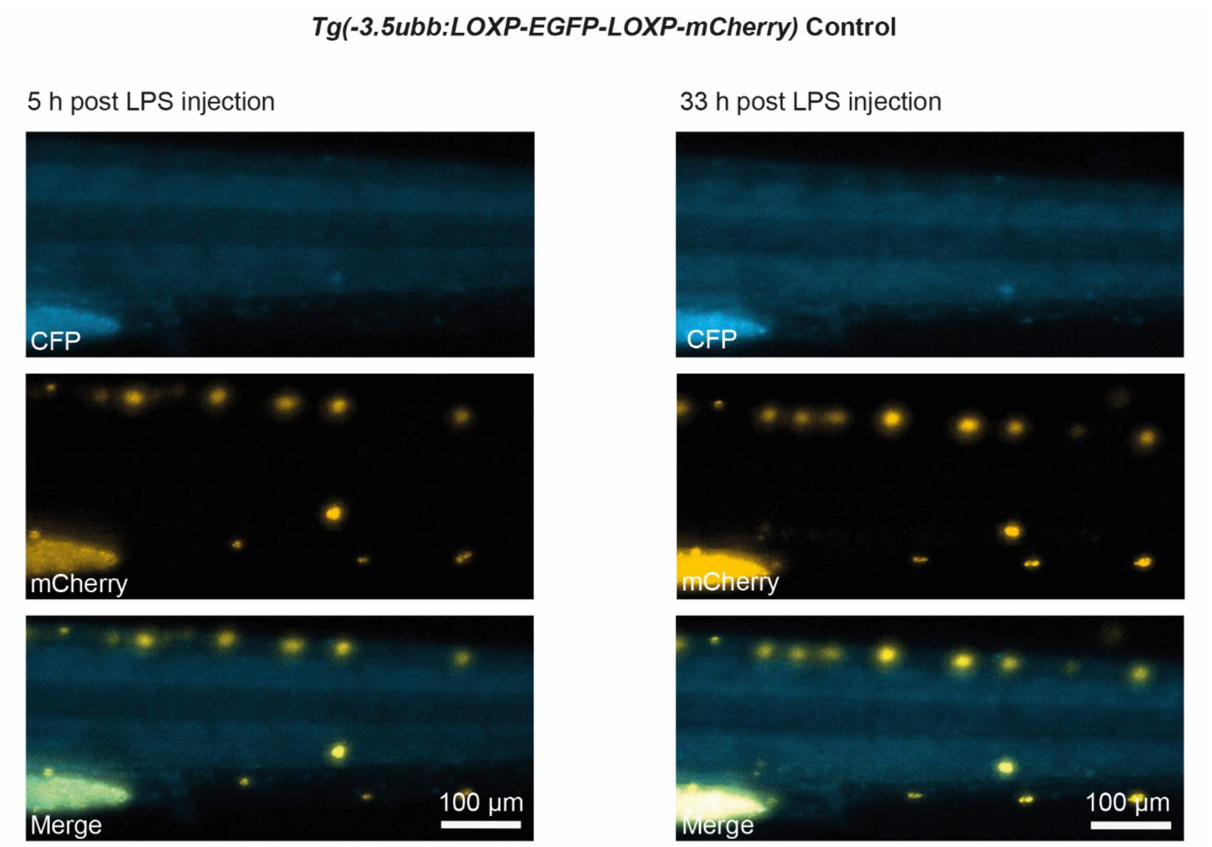


**Supplementary Figure 3: Control images for testing functionality of iCre**
As a control for testing of *Tg(irg1:icre-p2A mTurquoise2)* in *Tg(-3.5ubb:LOXP-EGFP-LOXP-mCherry)* animals in Fig.4 and Movie S1,*Tg(-3.5ubb:LOXP-EGFP-LOXP-mCherry)* control larvae were injected with LPS 2 dpf and screened. Images are shown in CFP, mCherry and merge 5 h (left) and 33 h (right) post LPS injection.

A

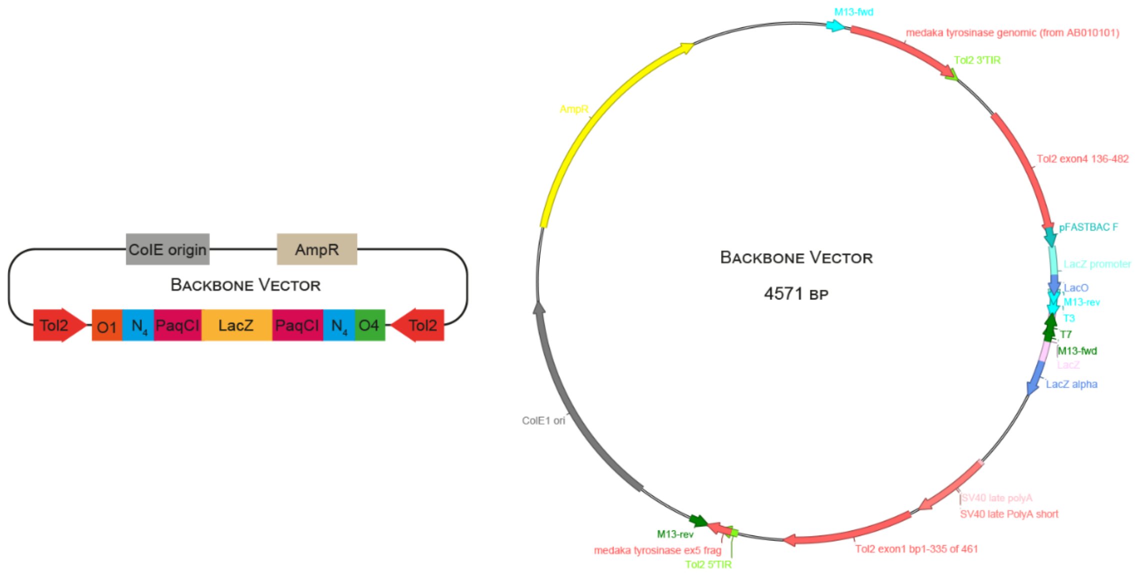


B

CCAGATACCTGAAACAAAACCCATCGTACGGCCAAGGAAGTCTCCAATAACTGTGATCCACCACAAGCGCCAGGGTTTTCCCAGTCACGACGTTGTAAAACGACG
GCCAGTCATGCATAATCCGCACGCATCTGGAATAAGGAAGTGCCATTCCGCCTGACCTCGCTGGCTAAGAACTCATCAGCCTCCCCGGTCCATCTACCCACG
TACCAATGCACCAATTGGCCACAATGACGGCTACTACATGGTGCCATTCCTTCCTCTTTATAGGAATGGAGACTACCTCCTGTCCAACAAGGCTCTTGGATAC
GAGTACGCCTACCTGTTGGACCCAGGTCATTGCACAACACCAGAAATGCCCTCTGATCTGCAAAAGACGTGAATATCTGTTCAGACACCCATATCCACTCTGT
TCCACACAGGTCAGAGGTTTGTCCAGGAGTTCTTGACAGAGGTGTAAAAAGTACTCAAAAATTTTACTCAAGTGAAAGTACAAGTACTTAGGGAAAATTTTACT
CAATTAAAAGTAAAAGTATCTGGCTAGAATCTTACTTGAGTAAAAGTAAAAAAGTACTCCATTAAAATTGTACTTGAGTATTAAGGAAGTAAAAGTAAAA
GCAAGAAAGAAAACTAGAGATTCTTGTTTAAGCTTTTAATCTCAAAAAACATTAAATGAAATGCATACAAGGTTTTATCCTGCTTTAGAACTGTTTGTATTTAA
TTATCAAACTATAAGACAGACAATCTAATGCCAGTACACGCTACTCAAAGTTGTAAAACCTCAGATTTAACTTCAGTAGAAGCTGATTCTCAAAATTGT
TAGTGTCAAGCCTAGCTCTTTTGGGGCTGAAAAGCAATCCTGCAGTGCTGAAAAGCCTCTCACAGGCAGCCGATGCGGGAAGAGGTGTATTAGTCTTGATAGA
GAGGCTGCAAATAGCAGGAAACGTGAGCAGAGACTCCCTGGTGTCTGAAACACAGGCCAGATGGGCCCTCGAGTCCGGATTATTCATACCGTCCCATTACGCCA
AGCTATTTCCTCTGGCAGGTGGCGCAACGCAATTAATGTGAGTTAGCTCACTCATTAGGCACCCCAGGCTTTACACTTTATGCTTCCGGCTCGTATGTTGTGTGG
AATTGTGAGCGGATAACAATTTCACACAGGAAACAGCTATGACCATGATTACGCCAAGCGCGCAATTAACCCTCACTAAAGGGAACAAAAGCTGGAGGACCTCG
AGGGGGGGCCCGGTACCCAATTCGCCCTATAGTGAGTCGTATTACGCGCGCTCACTGGCCGTCGTTTTACAACGTCGTGACTGGGAAAACCCTGGCGTTACC
CAACTTAATCGCCTTGCAGCACATCCCCCTTTCGCCAGCTGGCGTAATAGCGAAGAGGCCCGCACCGATCGCCCTTCCCAACAGTTGCGCAGCCTGAATGGCGAAT
GGCACCTGCTTCCAGGAATATTCTCAGTATCCCCTGCTTATGCACTCACCACTTTTTCCCATTATTTATTCATGTTGTGTTGCTAAATTCAACTACATTCGTATT
ACCATGTTGTTTCATTTGTTTTTTTTTATCTCAGCTGTCAAAATAAAAGCATTGAAATACAACATTGTTCTATTTTTCATTGTGGTTTTGCATGCCCGGTAAA
TAAAAAAACAAGGTAATAAAAATAAGATGCGAAATGAGAAGTATGATGATGATCCAGACATGATAAGATACATTGATGAGTTTGGACAAACCACAACTAGAA
TGCAGTGAAAAAAATGCTTTATTTGTGAAATTTGTGATGCTATTGCTTTATTTGTAACCATTATAAGCTGCAATAAACAAGTTAACAACAACAATTGCAT
TCATTTTATGTTTCAGGTTCAGGGGGAGGTGTGGGAGGTTTTTTAAAGCAAGTAAAACCTCTACAAATGTGGTATGGCTGATTATGATCCTCTAGATCAGATC
TGCGAAGATACGGCCACGGGTGCTCTTGATCCTGTGGCTGATTTTGGACTGTGCTGCTCGCAGCTGCTGATGAATCACATACTTCCTCCATTTTCTTCCACTGATT
GACTGTTATAATTTCCCTAATTTCCAGGTCAAGGTGCTGTGCATTGTGGTAATAGATGTGACATGACGTCACTTCCAAAGGACCAATGAACATGTCTGACCAATTT
CATATAATGTGAAAACGATTTTCATAGGCAGAATAAATAACATTTAAATTAAACTGGGCATCAGCGCAATTCAATTGGTTTGGTAATAGCAAGGGAAAATAG
AATGAAGTGATCTCCAAAAAATAAGTACTTTTTGACTGTAAATAAAATTGTAAGGAGTAAAAAGTACTTTTTTTTCTAAAAAAATGTAATTAAGTAAAAG
TAAAAGTATTGATTTTTAATTGTACTCAAGTAAAGTAAAAATCCCCAAAAATAATACTTAAGTACAGTAATCAAGTAAAATTACTCAAGTACTTTACACCTC
TGGTTCTTGACCCCCTACCTTCAGCAAGCCCAGCAGATCAGGCTAGGTGGAGGCTCAGTGATGATAAGTCTGCGATGGTGGATGCATGTGTCATGGTCATAGCT
GTTTCCTGTGTGAAATTGTTATCCGCTCAGAGGGCACAATCCTATTCCGCGCTATCCGACAATCTCCAAGACATTAGGTGGAGTTCAGTTCGGCGTATGGCATA
TGTCGCTGGAAAGAACATGTGAGCAAAAGGCCAGCAAAAGGCCAGGAACCGTAAAAAGGCCGCGTTGCTGGCGTT
TTTCCATAGGCTCCGCCCCCCTGACGAGCATCACAAAAATCGACGCTCAAGTCAGAGGTGGCGAAACCCGACAGGACTATAAAGATACCAGGCGTTTCCCCCTGG
AAGCTCCCTCGTGCGCTCTCCTGTTCCGACCCTGCCGCTTACCGGATACCTGTCCGCCTTTCTCCCTTCGGGAAGCGTGGCGCTTTCTCATAGCTCACGCTG
TAGGTATCTCAGTTCGGTGTAGGTCGTTCGCTCCAAGCTGGGCTGTGTGCACGAACCCCCCGTTCAGCCCGACCGCTGCGCCTTATCCGGTAACTATCGTCTT
GAGTCCAACCCGGTAAGACACGACTTATCGCCACTGGCAGCAGCCACTGGTAACAGGATTAGCAGAGCGAGGTATGTAGGCGGTGCTACAGAGTTC
TTGAAGTGGTGGCCTAACTACGGCTACACTAGAAGAACAGTATTTGGTATCTGCGCTCTGCTGAAGCCAGTTACCTTCGGAAAAAGAGTTGGTAGCTCTTGATC
CGGCAAACAAACCACCGCTGGTAGCGGTGGTTTTTTTGTTTGCAAGCAGCAGATTACGCGCAGAAAAAAAGGATCTCAAGAAGATCCTTTGATCTTTTC
TACGGGGTCTGACGCTCTATTCAACAAAGCCGCCGTCCCGTCAAGTCAGCGTAAATGGGTAGGGGGCTTCAAATCGTCCTCGTGATACCAATTCGGAGCCTGCTT
TTTTGTACAAACTTGTTGATAATGGCAATTCAAGGATCTTCACCTAGATCCTTTTAAATTAAAAATGAAGTTTTAAATCAATCTAAAGTATATATGAGTAAACT
TGGTCTGACAGTTACCAATGCTTAATCAGTGAGGCACCTATCTCAGCGATCTGTCTATTTCGTTCATCCATAGTTGCCTGACTCCCCGTCGTGTAGATAACTAC
GATACGGGAGGGCTTACCATCTGGCCCCAGTGCTGCAATGATACCGCGAGAGCCACGCTCACCGGCTCCAGATTTATCAGCAATAAACCAGCCAGCCGGAAGG
GCCGAGCGCAGAAGTGGTCCTGCAACTTTATCCGCCTCCATCCAGTCTATTAATTGTTGCCGGGAAGCTAGAGTAAGTAGTTCGCCAGTTAATAGTTTGCGCAAC
GTTGTTGCCATTGCTACAGGCATCGTGGTGTCACGCTCGTCGTTTGGTATGGCTTCATTCAGCTCCGGTTCCCAACGATCAAGGCGAGTTACATGATCC
CCCATGTTGTGCAAAAAAGCGGTTAGCTCCTTCGGTCCTCCGATCGTTGTCAGAAGTAAGTTGGCCGCAGTGTTATCACTCATGGTTATGGCAGCACTGCA
TAATTCTCTTACTGTCATGCCATCCGTAAGATGCTTTTCTGTGACTGGTGAGTACTCAACCAAGTCATTCTGAGAATAGTGTATGCGGCGACCGAGTTGCTCTT
GCCCGGCGTCAATACGGGATAATACCGCGCCACATAGCAGAACTTTAAAAGTGCTCATCATTGGAAAACGTTCTTCGGGGCGAAAACTCTCAAGGATCTTACCGC
TGTTGAGATCCAGTTCGATGTAACCCACTCGTGCACCCAACTGATCTTCAGCATCTTTTACTTTCACCAGCGTTTCTGGGTGAGCAAAAACAGGAAGGCAAAAT
GCCGCAAAAAAGGGAATAAGGGCGACACGGAAATGTTGAATACTCATACTCTTCCTTTTTCAATATTATTGAAGCATTTATCAGGGTTATTGTCTCATGAGCG
GATACATATTTGAATGTATTTAGAAAAATAAACAAATAGGGGTTCCGCGCACATTTCCCCGAAAAGTG

**Supplementary Figure 4: Sequence of the backbone vector**

A: Schematic of the backbone vector used for all Golden Gate assemblies conducted in this study. The backbone vector consists of an ampicillin resistance (AmpR) and *E. coli* origin (ColE), *Tol2* sites, PaqCI recognition sites (PaqCI) along with 4-base spacer sequences ($N_4$) and two overhangs (O1, O4). B: Complete DNA sequence of the backbone vector. Different parts of the vector are indicated by color corresponding to the schematic in A.

**A** *runx1+23*

TTTCCCCACCTGCCCTTTTCCGGGGTGGGAGGTGTAAGTTCCACCCCCACCCTTCCTGACACGCTCCT
GAACCCTGGCCACTGCACCTGGCTAGGTTCTCACTTCTCTGGGAAGCATCTAGAAACAGGACCTCTCA
CCCACCCCTCCCGGCTGGGCTCTAGGGTGGGGCCCTCACTACCTCTTTTCTTCTCAAAGAGCCTGGGA
TGCTGACAGCCTCAGATGGAGGCATCCTGTTTGTCGAAAAATAAACCGGCAGTTGAAGCCGGGTTGCA
AGAGCGAGAAAACCGCAGGCCTGCGCGCCACTGATAACGTGGGCAGCTTGCTTTTGCAGCAGTTCCTA
GCTGCAGCGGCCCTGTGAAGGCCTGTGTCACCGCCTCCCTTCCTGTCTCCTCCTCACACCATCCCTCC
ATCGCTCCTTGCTGGCTCTACCAGCCACTTGCTGGACCCTTCAGCCACTGGGACCATTGCTTTCCATA
AAAAACTCCTTAGTGCAGTGCAGAGCTGATCAGAGGGTAGCAGGGAACCCTGACGCCTGATGGGTTGC
TGACACCTGAGATCCACTAGTCCAATCTGCTCAGAGAGGACAGAGTGGGCAGGAGCCAGCATTGGGTA
TATAAAGCTGAGCAGGGTCAGTTGCTTCTTACGTTTGCTTCTGATTCTGTTGTGTTGACTTGCAACCT
CAGAAACAGACATCCAAACCTTGCAGGTGGGGTTT

**B** *tdStayGold*

TTTCCCCACCTGCCCTTCAAACATGGGAGAGGAGCTGTTCACAGGAGTGGTGATGGCTAGTACACCAT
TTAAATTTCAACTTAAAGGAACCATCAATGGCAAATCGTTTACCGTTGAAGGCGAAGGTGAAGGAAAT
TCACATGAAGGTTCTCATAAAGGAAAATATGTTTGTACAAGTGGAAAACTACCGATGTCATGGGCAGC
ACTTGGAACATCCTTTGGTTATGGAATGAAATATTATACCAAATATCCTAGTGGACTGAAGAACTGGT
TTCATGAAGTAATGCCTGAAGGCTTTACCTACGATCGTCATATTCAATATAAAGGCGATGGGAGTATC
CATGCAAAACACCAACACTTTATGAAAAATGGACTTATCACAACATTGTTGAATTTACTGGTCAGGA
TTTTAAAGAAAATAGTCCAGTCTTAACTGGAGATATGAATGTCTCATTACCGAATGAAGTTCAACATA
TACCCAGAGATGATGGAGTAGAATGCCCAGTGACCTTGCTTTATCCTTTATTATCGGATAAATCAAAA
TGCGTTGAGGCTCACCAAAATACAATCTGCAAGCCTCTTCATAATCAACCAGCACCTGATGTCCCATA
TCACTGGATTCGTAAACAATACACACAAAGCAAAGATGATACCGAGGAACGTGATCATATTTGTCAAT
CAGAGACTCTCGAAGCACACTTAGGAAACCCTTGGCACGAGCCTAGCGCTAGCGCTGTGAGTGCCGGC
GGTTCTGCAGGCGGGTCTGCAGGAGGTTCAGCTGGTGGTAGTGCAGGCGGCTCTGCAGGCGGTGGGGA
AGAACTGTTTACCGGCGTTGTTATGGCATCCACTCCTTTCAAGTTCCAGCTCAAGGGTACAATAAACG
GAAAGTCCTTCACAGTAGAGGGTGAAGGCGAGGGTAACAGTCACGAGGGGTCCCACAAGGGCAAGTAC
GTCTGCACCAGCGGTAAGCTGCCAATGTCTTGGGCTGCGCTGGGTACGAGTTTCGGGTACGGATGAA
GTACTACACAAAGTACCCATCTGGCCTTAAGAATTGGTTCCACGAGGTCATGCCAGAGGGGTTCACGT
ATGACAGGCACATCCAGTACAAAGGGGACGGCTCAATACACGCGAAGCATCAGCATTTCATGAAGAAC
GGTACATACCATAATATAGTAGAGTTCACAGGACAAGACTTCAAGGAGAACAGCCCTGTGCTAACAGG
GGACATGAACGTATCCCTGCCTAACGAGGTACAGCACATCCCTCGAGACGACGGCGTGGAGTGTCCCG
TTACGCTACTGTACCCACTGCTTTCCGACAAGTCTAAGTGTGTGGAAGCCCATCAGAACACGATTTGT
AAACCCCTGCACAACCAGCCCGCCCCCGACGTGCCCTACCATTGGATCCGAAAGCAGTATACCCAGTC
TAAGGACGACACTGAAGAGCGCGACCACATCTGCCAGAGCGAAACACTAGAGGCCCATTTGGGCAATC
CCTGGCATGAACCGTCCGCCAGTGCAGTCTAGGCTACCTTGCAGGTGGGGTTT

**C** *LNGFR*

TTTCCCCACCTGCCCTTCAAACATGGGAGCTGGAGCTACAGGCAGAGCTATGGATGGACCTAGACTGC
TGCTGCTGCTGCTGCTGGGTGTGAGCCTGGGAGGAGCTAAGGAAGCTTGTCCCACAGGACTGTATACT
CACTCTGGAGAGTGTTGTAAGGCCTGTAACCTGGGAGAGGGCGTGGCTCAGCCATGTGGAGCTAATCA
GACAGTGTGCGAGCCCTGTCTGGACTCTGTGACTTTCAGTGACGTGGTGAGTGCTACCGAGCCCTGTA
AACCATGCACCGAGTGCGTGGGACTGCAGTCTATGAGTGCTCCATGCGTGGAGGCTGACGACGCCGTG
TGCAGATGCGCTTACGGATATTACCAGGACGAGACAACAGGAAGATGCGAGGCTTGTAGAGTCTGCGA
AGCAGGAAGCGGACTGGTGTTCAGCTGTCAAGATAAACAGAACACAGTGTGCGAGGAGTGCCCAGACG
GAACCTATAGCGACGAGGCCAACCATGTCGATCCTTGCCTCCCTTGTACAGTGTGCGAGGACACTGAG
AGACAGCTGCGAGAGTGTACCAGATGGGCAGACGCTGAGTGTGAGGAGATTCCAGGAAGATGGATCAC
AAGAAGTACACCTCCAGAGGGCTCCGACAGCACAGCCCCATCCACACAGGAGCCTGAAGCTCCACCTG

AGCAGGACCTGATCGCTTCCACTGTGGCAGGAGTCGTGACAACTGTGATGGGCAGCTCTCAGCCCGTGGTGACACGTGGCACAACCGACAACCTGATCCCAGTGTATTGCTCCATCCTGGCCGCTGTGGTGGTGGGCCTGGTGGCTTACATCGCTTTCAAACGCTGAGCTACCTTGCAGGTGGGGTT

**D** *rac2*$^{WT}$

ACTGACTGCACCTGCCCTTCTCGATGCAAGCAATAAAGTGTGTGGTGGTCGGAGATGGAGCTGTGGGAAAGACCTGTCTTCTCATCAGCTACACTACCAATGCGTTCCCCGGGGAGTACATTCCCACAGTGTTTGATAACTACTCTGCAAATGTAATGGTGGATAGCAAACCAGTCAACCTGGGACTCTGGGATACAGCCGGACAGGAAGATTATGACAGACTGCGGCCACTCTCCTACCCGCAGACGGATGTGTTTCTTATCTGTTTCTCTTTGGTGAGCCCAGCATCATTCGAAAATGTCAGAGCCAAGTGGTACCCAGAGGTGAGGCATCACTGCCCTTCCACTCCAATTATCCTGGTTGGCACCAAGCTTGACTTGAGAGATGAGAAGGAGACCATCGAGAAGCTGAAGGAGAAGAAACTGGCACCGATCACTTACCCACAGGGTCTCGCATTGGCCAAAGAAATAGATGCAGTAAAATACCTGGAGTGTTCGGCCCTCACTCAGAGAGGGCTAAAAACAGTGTTTGATGAGGCGATTCGCGCTGTGCTCTGCCCACAGCCCACCAAGGTCAAGAAGAAGGGCTGCGTGATGCTCTAAAGGACCTTGCAGGTGACTGACTG

**E** *rac2*$^{D57N}$

ACTGACTGCACCTGCCCTTCTCGATGCAAGCAATAAAGTGTGTGGTGGTCGGAGATGGAGCTGTGGGAAAGACCTGTCTTCTCATCAGCTACACTACCAATGCGTTCCCCGGGGAGTACATTCCCACAGTGTTTGATAACTACTCTGCAAATGTAATGGTGGATAGCAAACCAGTCAACCTGGGACTCTGGAATACAGCCGGACAGGAAGATTATGACAGACTGCGGCCACTCTCCTACCCGCAGACGGATGTGTTTCTTATCTGTTTCTCTTTGGTGAGCCCAGCATCATTCGAAAATGTCAGAGCCAAGTGGTACCCAGAGGTGAGGCATCACTGCCCTTCCACTCCAATTATCCTGGTTGGCACCAAGCTTGACTTGAGAGATGAGAAGGAGACCATCGAGAAGCTGAAGGAGAAGAAACTGGCACCGATCACTTACCCACAGGGTCTCGCATTGGCCAAAGAAATAGATGCAGTAAAATACCTGGAGTGTTCGGCCCTCACTCAGAGAGGGCTAAAAACAGTGTTTGATGAGGCGATTCGCGCTGTGCTCTGCCCACAGCCCACCAAGGTCAAGAAGAAGGGCTGCGTGATGCTCTAAAGGACCTTGCAGGTGACTGACTG

**F** polyA-gRNA (Control):U6

AGCGCCCAATACGCAAACCGCCTCTCCCCGCGCGTTGGCCGATTCATTAATGCAGCTGGCACGACAGGTTTCCCGACTGGAAAGCGGGCAGTGAGCGCAACGCAATTAATGTGAGTTAGCTCACTCATTAGGCACCCCAGGCTTTACACTTTATGCTTCCGGCTCGTATGTTGTGTGGAATTGTGAGCGGATAACAATTTCACACAGGAAACAGCTATGACCATGATTACGCCAAGCTATTTAGGTGACACTATAGAATACTCAAGCTATGCATCAAGCTTGGTACCGAGCTCGGATCCACTAGTAACGGCCGCCAGTGTGCTGGAATTCGCCCTTCACCTGCCCTTCGTAAGATCTATAATTCACTGGCCGTCGTTTTACGGTACCATCGATGATGATCCAGACATGATAAGATACATTGATGAGTTTGGACAAACCACAACTAGAATGCAGTGAAAAAAATGCTTTATTTGTGAAATTTGTGATGCTATTGCTTTATTTGTAACCATTATAAGCTGCAATAAACAAGTTAACAACAACAATTGCATTCATTTTATGTTTCAGGTTCAGGGGGAGGTGTGGGAGGTTTTTTAAAGCAAGTAAAACCTCTACAAATGTGGTATGGCTGATTATGATCCTCTAGATCGAGGTCTCTGACTAAAAAAGCACCGACTCGGTGCCACTTTTTCAAGTTGATAACGGACTAGCCTTATTTAAACTTGCTATGCTGTTTCCAGCATAGCTCTTAAACAGAGACGGTCGACAGTCTGCAGTGTCGTCTCTCGAACCAAGAGCTGGAGGGAGAGCTATATATACCAGGGACTTCTGGGTATGTTTTTGGGAGGTGGTGAGTGACTAAACCACTTATTCAGCTCCCTTAGATCAAGTCTGACCCTATATCATGGTGACATAAACCTGCAAACTGATAAAACCTGAAGGATCTCAAATCCAGAGTTTGTGTGAGGGATTACCGTGGTAATTTTCAAACTGTAAAGCATATGCAAAATTATCTGGTCTTGGCTTGAGTGATTGGGTGTCTCGGTGTGATGCAGGGACGTTTTCAGTGACGTGTCCTTCTCCCTCCCCCACAGGCATGCGCAGAACATTTCCCCCCTCCTTGAAGACCAGAACAAAAGACGCCGAGAGCAGGAAACTCGTCTTACTGAATGACCGAGGCTGGAGAAAGTCGACCCTAGGAGGACCTTGCAGGTGAAGGGCGAATTCTGCAGATATCCATCACACTGGCGGCCGCTCGAGCATGCATCTAGAGGGCCCAATTCGCCCTATAGTGAGTCGTATTACAATTCACTGGCCGTCGTTTTACAACGTCGTGACTGGGAAAACCCTGGCGTTACCCAACTTAATCGCCTTGCAGCACATCCCCCTTTCGCCAGCTGGCGTAATAGCGAAGAGGCCCGCACCGATCGCCCTTCCCAACAGTTGCGCAGCCTATACGTACGGCAGTTTAAGGTTTACACCTATAAAAGAGAGAGCCGTTATCGTCTGTTTGTGGATGTACAGAGTGATATTATTGACACGCCGGGGCGACGGATGGTGATCCCCCTGGCCAGTGCACGTCTGCTGTCAGATAAAGTCTCCCGTGAACTTTACCCGGTGGTGCATATCGGGGATGAAAGCTGGCGCATGATGACCACCGATATGGCCAGTGTGCCGGTCTCCGTTATCGGGGAAGAAGTGGCTGATCTCAGCCACCGCGAAAATGACATCAAAAACGCCATTAACCTGATGTTCTGGGGAATATAAATGTCAGGCATGAGATTATCAAAAAGGATCTTCACCTAGATCCTTTTCACGTAGAAAGCCAGTCCGCAGAAACGGTGCTGACCCCGGATGAATGTCAGCTACTGGGCTATCTGGACAAGGGAAAACGCAAGCGCAAAGAGAAAGCAGGTAGCTTGCAGTGGGCTTACATGGCGATAGCTAGACTGG

GCGGTTTTATGGACAGCAAGCGAACCGGAATTGCCAGCTGGGGCGCCCTCTGGTAAGGTTGGGAAGCCCTGCAAA
GTAAACTGGATGGCTTTCTCGCCGCCAAGGATCTGATGGCGCAGGGGATCAAGCTCTGATCAAGAGACAGGATGA
GGATCGTTTCGCATGATTGAACAAGATGGATTGCACGCAGGTTCTCCGGCCGCTTGGGTGGAGAGGCTATTCGGC
TATGACTGGGCACAACAGACAATCGGCTGCTCTGATGCCGCCGTGTTCCGGCTGTCAGCGCAGGGGCGCCCGGTT
CTTTTTGTCAAGACCGACCTGTCCGGTGCCCTGAATGAACTGCAAGACGAGGCAGCGCGGCTATCGTGGCTGGCC
ACGACGGGCGTTCCTTGCGCAGCTGTGCTCGACGTTGTCACTGAAGCGGGAAGGGACTGGCTGCTATTGGGCGAA
GTGCCGGGGCAGGATCTCCTGTCATCTCACCTTGCTCCTGCCGAGAAAGTATCCATCATGGCTGATGCAATGCGG
CGGCTGCATACGCTTGATCCGGCTACCTGCCCATTCGACCACCAAGCGAAACATCGCATCGAGCGAGCACGTACT
CGGATGGAAGCCGGTCTTGTCGATCAGGATGATCTGGACGAAGAGCATCAGGGGCTCGCGCCAGCCGAACTGTTC
GCCAGGCTCAAGGCGAGCATGCCCGACGGCGAGGATCTCGTCGTGACCCATGGCGATGCCTGCTTGCCGAATATC
ATGGTGGAAAATGGCCGCTTTTCTGGATTCATCGACTGTGGCCGGCTGGGTGTGGCGGACCGCTATCAGGACATA
GCGTTGGCTACCCGTGATATTGCTGAAGAGCTTGGCGGCGAATGGGCTGACCGCTTCCTCGTGCTTTACGGTATC
GCCGCTCCCGATTCGCAGCGCATCGCCTTCTATCGCCTTCTTGACGAGTTCTTCTGAATTATTAACGCTTACAAT
TTCCTGATGCGGTATTTTCTCCTTACGCATCTGTGCGGTATTTCACACCGCATACAGGTGGCACTTTTCGGGGAA
ATGTGCGCGGAACCCCTATTTGTTTATTTTTCTAAATACATTCAAATATGTATCCGCTCATGAGACAATAACCCT
GATAAATGCTTCAATAATAGCACGTGAGGAGGGCCACCATGGCCAAGTTGACCAGTGCCGTTCCGGTGCTCACCG
CGCGCGACGTCGCCGGAGCGGTCGAGTTCTGGACCGACCGGCTCGGGTTCTCCCGGGACTTCGTGGAGGACGACT
TCGCCGGTGTGGTCCGGGACGACGTGACCCTGTTCATCAGCGCGGTCCAGGACCAGGTGGTGCCGGACAACACCC
TGGCCTGGGTGTGGGTGCGCGGCCTGGACGAGCTGTACGCCGAGTGGTCGGAGGTCGTGTCCACGAACTTCCGGG
ACGCCTCCGGGCCGGCCATGACCGAGATCGGCGAGCAGCCGTGGGGGCGGGAGTTCGCCCTGCGCGACCCGGCCG
GCAACTGCGTGCACTTCGTGGCCGAGGAGCAGGACTGACACGTGCTAAAACTTCATTTTTAATTTAAAAGGATCT
AGGTGAAGATCCTTTTTGATAATCTCATGACCAAAATCCCTTAACGTGAGTTTTCGTTCCACTGAGCGTCAGACC
CCGTAGAAAAGATCAAAGGATCTTCTTGAGATCCTTTTTTTCTGCGCGTAATCTGCTGCTTGCAAACAAAAAAAC
CACCGCTACCAGCGGTGGTTTGTTTGCCGGATCAAGAGCTACCAACTCTTTTTCCGAAGGTAACTGGCTTCAGCA
GAGCGCAGATACCAAATACTGTCCTTCTAGTGTAGCCGTAGTTAGGCCACCACTTCAAGAACTCTGTAGCACCGC
CTACATACCTCGCTCTGCTAATCCTGTTACCAGTGGCTGCTGCCAGTGGCGATAAGTCGTGTCTTACCGGGTTGG
ACTCAAGACGATAGTTACCGGATAAGGCGCAGCGGTCGGGCTGAACGGGGGGTTCGTGCACACAGCCCAGCTTGG
AGCGAACGACCTACACCGAACTGAGATACCTACAGCGTGAGCTATGAGAAAGCGCCACGCTTCCCGAAGGGAGAA
AGGCGGACAGGTATCCGGTAAGCGGCAGGGTCGGAACAGGAGAGCGCACGAGGGAGCTTCCAGGGGGAAACGCCT
GGTATCTTTATAGTCCTGTCGGGTTTCGCCACCTCTGACTTGAGCGTCGATTTTTGTGATGCTCGTCAGGGGGGC
GGAGCCTATGGAAAAACGCCAGCAACGCGGCCTTTTTACGGTTCCTGGGCTTTTGCTGGCCTTTTGCTCACATGT
TCTTTCCTGCGTTATCCCCTGATTCTGTGGATAACCGTATTACCGCCTTTGAGTGAGCTGATACCGCTCGCCGCA
GCCGAACGACCGAGCGCAGCGAGTCAGTGAGCGAGGAAGCGGAAG

**G** polyA-gRNA (GFP):U6

AGCGCCCAATACGCAAACCGCCTCTCCCCGCGCGTTGGCCGATTCATTAATGCAGCTGGCACGACAGGTTTCCCG
ACTGGAAAGCGGGCAGTGAGCGCAACGCAATTAATGTGAGTTAGCTCACTCATTAGGCACCCCAGGCTTTACACT
TTATGCTTCCGGCTCGTATGTTGTGTGGAATTGTGAGCGGATAACAATTTCACACAGGAAACAGCTATGACCATG
ATTACGCCAAGCTATTTAGGTGACACTATAGAATACTCAAGCTATGCATCAAGCTTGGTACCGAGCTCGGATCCA
CTAGTAACGGCCGCCAGTGTGCTGGAATTCGCCCTTCACCTGCCCTTCGTAAGATCTATAATTCACTGGCCGTCG
TTTTACGGTACCATCGATGATGATCCAGACATGATAAGATACATTGATGAGTTTGGACAAACCACAACTAGAATG
CAGTGAAAAAAATGCTTTATTTGTGAAATTTGTGATGCTATTGCTTTATTTGTAACCATTATAAGCTGCAATAAA
CAAGTTAACAACAACAATTGCATTCATTTTATGTTTCAGGTTCAGGGGGAGGTGTGGGAGGTTTTTTAAAGCAAG
TAAAACCTCTACAAATGTGGTATGGCTGATTATGATCCTCTAGATCGAGGTCTCTGACTAAAAAAGCACCGACTC
GGTGCCACTTTTTCAAGTTGATAACGGACTAGCCTTATTTAAACTTGCTATGCTGTTTCCAGCATAGCTCTTAAA
CCCGTAGGTGGCATCGCCCTCGCCGAACCAAGAGCTGGAGGGAGAGCTATATATACCAGGGACTTCTGGGTATGT
TTTTGGGAGGTGGTGAGTGACTAAACCACTTATTCAGCTCCCTTAGATCAAGTCTGACCCTATATCATGGTGACA
TAAACCTGCAAACTGATAAAACCTGAAGGATCTCAAATCCAGAGTTTGTGTGAGGGATTACCGTGGTAATTTTCA
AACTGTAAAGCATATGCAAAATTATCTGGTCTTGGCTTGAGTGATTGGGTGTCTCGGTGTGATGCAGGGACGTTT
TCAGTGACGTGTCCTTCTCCCTCCCCCACAGGCATGCGCAGAACATTTCCCCCCTCCTTGAAGACCAGAACAAAA
GACGCCGAGAGCAGGAAACTCGTCTTACTGAATGACCGAGGCTGGAGAAAGTCGACCCTAGGAGGACCTTGCAGG
TGAAGGGCGAATTCTGCAGATATCCATCACACTGGCGGCCGCTCGAGCATGCATCTAGAGGGCCCAATTCGCCCT
ATAGTGAGTCGTATTACAATTCACTGGCCGTCGTTTTACAACGTCGTGACTGGGAAAACCCTGGCGTTACCCAAC
TTAATCGCCTTGCAGCACATCCCCCTTTCGCCAGCTGGCGTAATAGCGAAGAGGCCCGCACCGATCGCCCTTCCC

AACAGTTGCGCAGCCTATACGTACGGCAGTTTAAGGTTTACACCTATAAAAGAGAGAGCCGTTATCGTCTGTTTG
TGGATGTACAGAGTGATATTATTGACACGCCGGGGCGACGGATGGTGATCCCCCTGGCCAGTGCACGTCTGCTGT
CAGATAAAGTCTCCCGTGAACTTTACCCGGTGGTGCATATCGGGGATGAAAGCTGGCGCATGATGACCACCGATA
TGGCCAGTGTGCCGGTCTCCGTTATCGGGGAAGAAGTGGCTGATCTCAGCCACCGCGAAAATGACATCAAAAACG
CCATTAACCTGATGTTCTGGGGAATATAAATGTCAGGCATGAGATTATCAAAAAGGATCTTCACCTAGATCCTTT
TCACGTAGAAAGCCAGTCCGCAGAAACGGTGCTGACCCCGGATGAATGTCAGCTACTGGGCTATCTGGACAAGGG
AAAACGCAAGCGCAAAGAGAAAGCAGGTAGCTTGCAGTGGGCTTACATGGCGATAGCTAGACTGGGCGGTTTTAT
GGACAGCAAGCGAACCGGAATTGCCAGCTGGGGCGCCCTCTGGTAAGGTTGGGAAGCCCTGCAAAGTAAACTGGA
TGGCTTTCTCGCCGCCAAGGATCTGATGGCGCAGGGGATCAAGCTCTGATCAAGAGACAGGATGAGGATCGTTTC
GCATGATTGAACAAGATGGATTGCACGCAGGTTCTCCGGCCGCTTGGGTGGAGAGGCTATTCGGCTATGACTGGG
CACAACAGACAATCGGCTGCTCTGATGCCGCCGTGTTCCGGCTGTCAGCGCAGGGGCGCCCGGTTCTTTTTGTCA
AGACCGACCTGTCCGGTGCCCTGAATGAACTGCAAGACGAGGCAGCGCGGCTATCGTGGCTGGCCACGACGGGCG
TTCCTTGCGCAGCTGTGCTCGACGTTGTCACTGAAGCGGGAAGGGACTGGCTGCTATTGGGCGAAGTGCCGGGGC
AGGATCTCCTGTCATCTCACCTTGCTCCTGCCGAGAAAGTATCCATCATGGCTGATGCAATGCGGCGGCTGCATA
CGCTTGATCCGGCTACCTGCCCATTCGACCACCAAGCGAAACATCGCATCGAGCGAGCACGTACTCGGATGGAAG
CCGGTCTTGTCGATCAGGATGATCTGGACGAAGAGCATCAGGGGCTCGCGCCAGCCGAACTGTTCGCCAGGCTCA
AGGCGAGCATGCCCGACGGCGAGGATCTCGTCGTGACCCATGGCGATGCCTGCTTGCCGAATATCATGGTGGAAA
ATGGCCGCTTTTCTGGATTCATCGACTGTGGCCGGCTGGGTGTGGCGGACCGCTATCAGGACATAGCGTTGGCTA
CCCGTGATATTGCTGAAGAGCTTGGCGGCGAATGGGCTGACCGCTTCCTCGTGCTTTACGGTATCGCCGCTCCCG
ATTCGCAGCGCATCGCCTTCTATCGCCTTCTTGACGAGTTCTTCTGAATTATTAACGCTTACAATTTCCTGATGC
GGTATTTTCTCCTTACGCATCTGTGCGGTATTTCACACCGCATACAGGTGGCACTTTTCGGGGAAATGTGCGCGG
AACCCCTATTTGTTTATTTTTCTAAATACATTCAAATATGTATCCGCTCATGAGACAATAACCCTGATAAATGCT
TCAATAATAGCACGTGAGGAGGGCCACCATGGCCAAGTTGACCAGTGCCGTTCCGGTGCTCACCGCGCGCGACGT
CGCCGGAGCGGTCGAGTTCTGGACCGACCGGCTCGGGTTCTCCCGGGACTTCGTGGAGGACGACTTCGCCGGTGT
GGTCCGGGACGACGTGACCCTGTTCATCAGCGCGGTCCAGGACCAGGTGGTGCCGGACAACACCCTGGCCTGGGT
GTGGGTGCGCGGCCTGGACGAGCTGTACGCCGAGTGGTCGGAGGTCGTGTCCACGAACTTCCGGGACGCCTCCGG
GCCGGCCATGACCGAGATCGGCGAGCAGCCGTGGGGGCGGGAGTTCGCCCTGCGCGACCCGGCCGGCAACTGCGT
GCACTTCGTGGCCGAGGAGCAGGACTGACACGTGCTAAAACTTCATTTTTAATTTAAAAGGATCTAGGTGAAGAT
CCTTTTTGATAATCTCATGACCAAAATCCCTTAACGTGAGTTTTCGTTCCACTGAGCGTCAGACCCCGTAGAAAA
GATCAAAGGATCTTCTTGAGATCCTTTTTTTCTGCGCGTAATCTGCTGCTTGCAAACAAAAAAACCACCGCTACC
AGCGGTGGTTTGTTTGCCGGATCAAGAGCTACCAACTCTTTTTCCGAAGGTAACTGGCTTCAGCAGAGCGCAGAT
ACCAAATACTGTCCTTCTAGTGTAGCCGTAGTTAGGCCACCACTTCAAGAACTCTGTAGCACCGCCTACATACCT
CGCTCTGCTAATCCTGTTACCAGTGGCTGCTGCCAGTGGCGATAAGTCGTGTCTTACCGGGTTGGACTCAAGACG
ATAGTTACCGGATAAGGCGCAGCGGTCGGGCTGAACGGGGGGTTCGTGCACACAGCCCAGCTTGGAGCGAACGAC
CTACACCGAACTGAGATACCTACAGCGTGAGCTATGAGAAAGCGCCACGCTTCCCGAAGGGAGAAAGGCGGACAG
GTATCCGGTAAGCGGCAGGGTCGGAACAGGAGAGCGCACGAGGGAGCTTCCAGGGGGAAACGCCTGGTATCTTTA
TAGTCCTGTCGGGTTTCGCCACCTCTGACTTGAGCGTCGATTTTTGTGATGCTCGTCAGGGGGGCGGAGCCTATG
GAAAAACGCCAGCAACGCGGCCTTTTTACGGTTCCTGGGCTTTTGCTGGCCTTTTGCTCACATGTTCTTTCCTGC
GTTATCCCCTGATTCTGTGGATAACCGTATTACCGCCTTTGAGTGAGCTGATACCGCTCGCCGCAGCCGAACGAC
CGAGCGCAGCGAGTCAGTGAGCGAGGAAGCGGAAG

**Supplementary Figure 5: Sequences of genes and constructs purchased as dsDNA fragments.** A: Sequence of *runx1+23* including PaqCI sites, O1 and O2. B: Sequence of *tdStayGold* including PaqCI sites, O2 and O3. C: Sequence of *LNGFR* including PaqCI sites, O2 and O3. D: Sequence of *rac2*$^{WT}$ including PaqCI sites, overhang sequence CTCG (O3B) and O4. E: Sequence of *rac2*$^{D57N}$ including PaqCI sites, O3B and O4. F: Sequence of 3E polyA-gRNA (Control):U6 including PaqCI sites, overhang sequence CGTA (O3C) and O4. G: Sequence of 3E polyA-gRNA (GFP):U6 including PaqCI sites, O3C and O4.

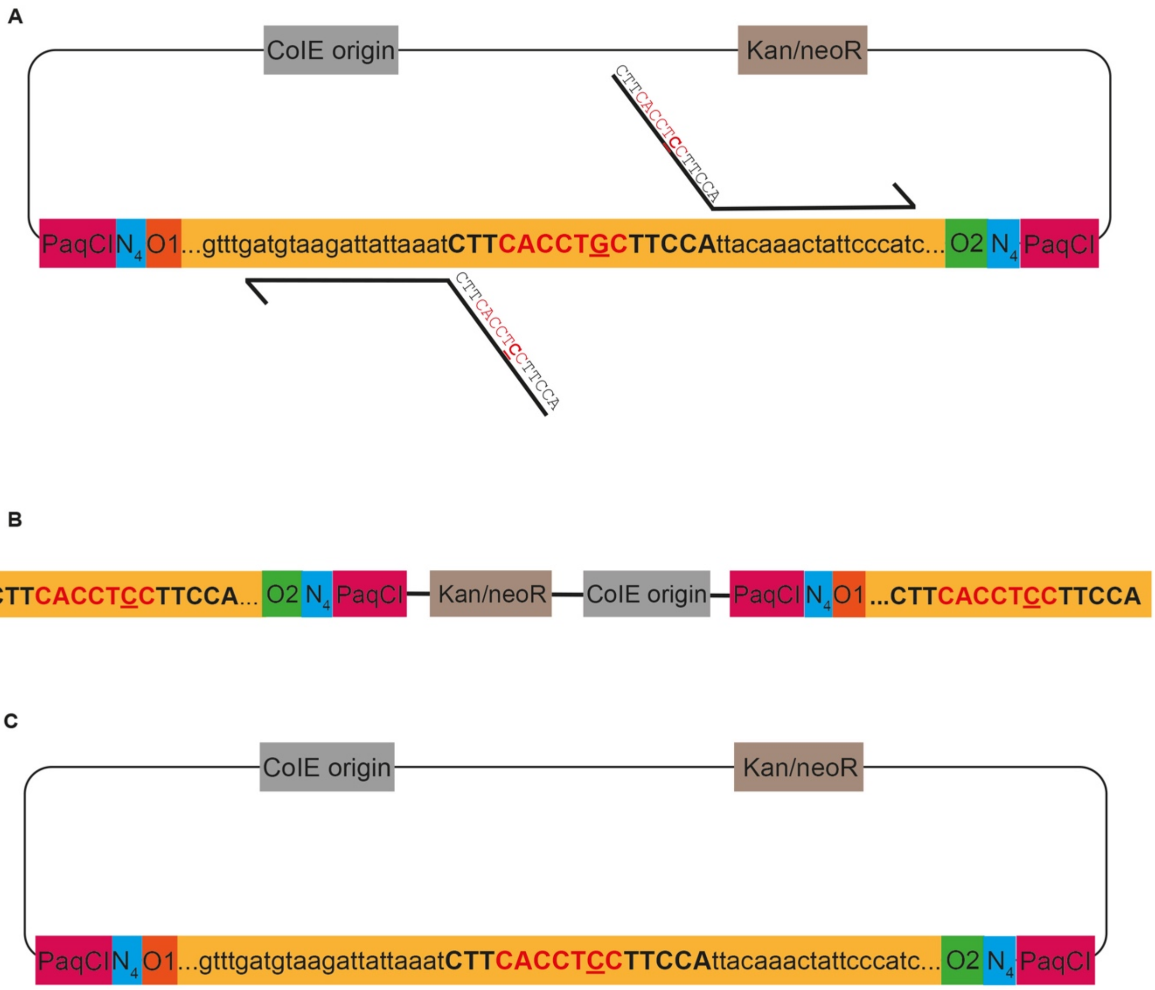


**Supplementary Figure 6: Domestication of endogenous PaqCI sites.**
A: Schematic of the 5E *ubb* promoter construct as an example of an insertion element containing an endogenous PaqCI site. The construct contains PaqCI sites (pink), spacer sequence ($N_4$, blue) and overhang sequences O1 (orange) and O2 (green). Primers were designed to introduce a single base mutation (highlighted in bold and underlined) in the endogenous PaqCI recognition site (red). B: PCR amplified *ubb* promoter with mutated endogenous PaqCI site and homology arms for In-Fusion cloning. C: In-Fusion reaction of the PCR product led to the final 5E *ubb* promoter construct with the mutated endogenous PaqCI site.

**Supplementary Table 1: Primer Sequences.**
PaqCI sites are shown in magenta, 4-base spacer sequences are shown in blue, overhang 1 is shown in orange, O1A in purple, O1B in brown O2 in green, O3 in pink, O3B in light green and O4 in turquoise. Bases mutated to eliminate endogenous PaqCI sites are highlighted in red.

| | Gene | Primer Direction | Primer Sequence 5' → 3' |
|---|---|---|---|
| 5' Element | *irg1* | Forward | TTTCCCCACCTGCTTCCTTCCGCAGTGAGGGGTATGGAAAA |
| | | Reverse | TTGTACCACCTGCTTCATTTGTGTGCGAGCTCTGAATCTTG |
| | *ubb* | Forward | TTTCCCCACCTGCCCTTTTCCTTCTAGAATTTGTCGAAACATTTATG |
| | | Reverse | AAACCCCACCTGCAAGGTTTGCTGTAAACAAATTCAAAGTAAGATTAG |
| | PaqCI mutation in *ubb* | Forward | CTTCACCT**C**CTTCCATTACAAACTATTCCCATC |
| | | Reverse | TGGAAGGAGGTGAAGATTTAATAATCTTACATCAAAC |
| | *ubb₁* | Forward | TTTCCCCACCTGCCCTTTTCCTTCTA |
| | | Reverse | AAACCCCACCTGCAAGGGCTATTGGTGAATCAGACTTGTGGAATCTG |
| | *ubb₂* | Forward | TTTCCCCACCTGCCCTTTAGCAAAGCGAATAAACAACCAAA |
| | | Reverse | AAACCCCACCTGCAAGGTCAAGTAAGAAAAATATCAAGCATATTAC |
| | *ubb₃* | Forward | TTTCCCCACCTGCCCTTTTGATCGGAAATAAGAAAAAAATATAAACGT |
| | | Reverse | ATTATACACCTGCAAGGTTTGCTGTAAA |
| | *mfap4* | Forward | TTTCCCCACCTGCCCTTTTCCGCGTTTCTTGGTACAGC |
| | | Reverse | AAACCCCACCTGCAAGGTTTGCACGATCTAAAGTCATGAAGAAAG |
| | *mpeg1.1* | Forward | (TTTCCC) CACCTGCTTCCTTCCTTGTTGGAGCACATCTGACA |
| | | Reverse | (TTTCCC) CACCTGCTTCATTTGTGTTTTGCTGTCTCCTGCAC |
| | *ubb:loxP GFP loxP* | Forward | TTTCCCCACCTGCCCTTTTCCACCAGCAAAGTTCTAGAATTTGTC |
| | | Reverse | AAACCCCACCTGCAAGGTTTGCGCCCTTTTTGGATCCA |
| | *hsp70* | Forward | TTTCCCCACCTGCCCTTTTCCTCAGGGGTGTCGC |
| | | Reverse | AAACCCCACCTGCAAGGTTTGCAGGAAAAAAAAACAATTAGAATTAATTTTATATT |
| | *lck* | Forward | TTTCCCCACCTGCCCTTTTCCGAATTCACAATCTCATCATCATCATCTGAGC |
| | | Reverse | AAACCCCACCTGCAAGGTTTGGTTTACTAAGCATGAGAGAAAATGGTGCAACTATA |
| | PaqCI Mutation in *lck* | Forward | GCA**GG**TCCTCGTCGCTCTGAGCAG |
| | | Reverse | CTCAGAGCGACGAGGACCTGCTGCAA |
| | *lyz* | Forward | TTTCCCCACCTGCCCTTTTCCCCTGATCACTGGTGTAGTGAACTC |
| | | Reverse | AAACCCCACCTGCAAGGTTTGTCGAGATTGTATCACTGCTGATATCTGC |
| | *runX1+23* | Forward | purchased from Twist Bioscience, containing PaqCI recognition site, O1 and O2 |
| | | Reverse | |
| Middle Element | *icre* | Forward | TTTCCCCACCTGCCCTTCAAACATG |
| | | Reverse | AAACCCCACCTGCAAGGTAGCGTCCCCATCCTCGAGCAGC |
| | *tdTomato* | Forward | TTTCCCCACCTGCCCTTCAAACATGGTGAGCAAGGGC |
| | | Reverse | AAACCCCACCTGCAAGGTAGCTTACTTGTACAGCTCGTCCA |
| | *tdTomato* (no stop) | Forward | (ACTGACTG) CACCTGCCCTTCAAACATGGTGAGCAAGGGC |
| | | Reverse | (CAGTCAGT) CACCTGCAAGGTAGCCTTGTACAGCTCGTCCATG |
| | *tdStayGold* | na | purchased from Twist Bioscience, containing PaqCI recognition site, O2 and O3 |
| | | na | |
| | *nitroreductase* | Forward | TTTCCCCACCTGCCCTTCAAACATGGCCTCCGGACTCA |
| | | Reverse | AAACCCCACCTGCAAGGTAGCCACTTCGGTTAAGGTGATGTTTTGC |
| | *tdTomato CAAX* | Forward | TTTCCCCACCTGCCCTTCAAACATGGTGAGCAAGGGCGAG |
| | | Reverse | AAACCCCACCTGCAAGGTAGCCTAGGAGAGCACACACTTGCA |
| | *mTurquoise2* | Forward | TTTCCCCACCTGCCCTTCAAACATGGTGAGCAAGGGC |
| | | Reverse | AAACCCCACCTGCAAGGTAGCTTACTTGTACAGCTCGTCCA |
| | *mNeonGreen* | Forward | TTTCCCCACCTGCCCTTCAAACATGGTGAGCAAGGGC |
| | | Reverse | AAACCCCACCTGCAAGGTAGCTTACTTGTACAGCTCGTCCA |
| | *mCitrine* | Forward | TTTCCCCACCTGCCCTTCAAACATGGTGAGCAAGGGC |
| | | Reverse | AAACCCCACCTGCAAGGTAGCTTACTTGTACAGCTCGTCCA |
| | *kid* | Forward | TTTCCCCACCTGCCCTTCAAAAGTCAGAATAGTGGACAG |
| | | Reverse | AAACCCCACCTGCAAGGTAGCCAGGAAGCGGAGCTAC |

| | | | |
|---|---|---|---|
| | *LNGFR* | Forward | purchased from Twist Bioscience, containing PaqCI recognition site, O2 and O3 |
| | | Reverse | |
| | *p2A* | Forward | (ACTGACTG)CACCTGCCCTTGCTACAGGAAGCGGAGCTACTAACTT |
| | | Reverse | (CAGTCAGT)CACCTGCAAGGCGAGCTAGGTCCAGGGTTCTCC |
| **3‘ Element** | *p2A tdTomato* | Forward | TTTCCCCACCTGCTTCCGCTACAGGAAGCGGAGCTACTAAC |
| | | Reverse | TTGTACCACCTGCTTCATCCTTTACTTGTACAGCTCGTCCA |
| | *p2A mTurquoise* | Forward | AAGGAACACCTGCAAAGGCTACAATGGTGAGCAAGGGC |
| | | Reverse | AAACCCCACCTGCAAGGTCCTTTACTTGTACAGCTCGTCCATG |
| | *ubb:polyA* | Forward | TTTCCCCACCTGCCCTTGCTACAATTCTCAGTATCCCCTGC |
| | | Reverse | AAACCCCACCTGCAAGGTCCTATACTTCTCATTTCGCATCTTATT |
| | *p2A dLanYFP* | Forward | TTTCCCCACCTGCCCTTGCTACAGGAAGCGGAGCTACTAACTT |
| | | Reverse | AAACCCCACCTGCAAGGTCCTTTACTTGTACAGCTCGTCCATG |
| | *mTurquoise2* | Forward | TTTCCCCACCTGCCCTTGCTACAGGAAGCGGAGCTACTAACTT |
| | | Reverse | AAACCCCACCTGCAAGGTCCTTTACTTGTACAGCTCGTCCATG |
| | *tdTomato* | Forward | AAGGAACACCTGCAAAGGCTACAATGGTGAGCAAGGGC |
| | | Reverse | AAACCCCACCTGCAAGGTCCTTTACTTGTACAGCTCGTCCATG |
| | *mNeonGreen* | Forward | AAGGAACACCTGCAAAGGCTACAATGGTGAGCAAGGGC |
| | | Reverse | AAACCCCACCTGCAAGGTCCTTTACTTGTACAGCTCGTCCATG |
| | *$rac2^{WT}$* | Forward | purchased from IDT, containing PaqCI recognition site, O3B and O4 |
| | | Reverse | |
| | *$rac2^{D57N}$* | Forward | purchased from IDT, containing PaqCI recognition site, O3B and O4 |
| | | Reverse | |
| | polyA-gRNA(Cntrl):U6 | Forward | purchased from IDT, containing PaqCI recognition site, O3C and O4 |
| | | Reverse | |
| | polyA-gRNA(GFP):U6 | Forward | purchased from IDT, containing PaqCI recognition site, O3C and O4 |
| | | Reverse | |

**Movie S1: Time-lapse imaging of *Tg(-3.5ubb:LOXP-EGFP-LOXP-mCherry)* injected with *Tg(irg1:icre-p2A mTurquoise2)* zebrafish for lineage tracing of individual macrophages.**

The *Tg(irg1:icre-p2A mTurquoise2)* construct was introduced into fertilized eggs from *Tg(-3.5ubb:LOXP-EGFP-LOXP-mCherry)* zebrafish at their single cell stage. Larvae were treated with PTU and LPS and time-lapse imaging was performed for 48h. The movie shows a macrophage-specific CFP and mCherry signal. Scale bar is 100 µm.

**Movie S2: Time-lapse imaging of *Tg(-3.5ubb:LOXP-EGFP-LOXP-mCherry)* control zebrafish.**
As a control to the Cre induced rearrangement of *Tg(-3.5ubb:LOXP-EGFP-LOXP-mCherry)* by *Tg(irg1:icre-p2A mTurquoise2)* in Fig.4 and Movie S1,*Tg(-3.5ubb:LOXP-EGFP-LOXP-mCherry)* control larvae were treated with PTU and LPS and time-lapse imaging was performed for 48h. The movie shows no macrophage-specific CFP and mCherry signal. Scale bar is 100 µm.
